# Genetic regulation of RNA splicing in human pancreatic islets

**DOI:** 10.1101/2021.11.11.468254

**Authors:** Goutham Atla, Silvia Bonas-Guarch, Anthony Beucher, Mirabai Cuenca-Ardura, Javier Garcia-Hurtado, Ignasi Moran, the T2DSystems consortium, Manuel Irimia, Rashmi B Prasad, Anna L. Gloyn, Lorella Marselli, Mara Suleiman, Thierry Berney, Eelco J P de Koning, Julie Kerr-Conte, Francois Pattou, Lorenzo Piemonti, Jorge Ferrer

## Abstract

Genetic variants that influence transcriptional regulation in pancreatic islets play a major role in the susceptibility to type 2 diabetes (T2D). For many susceptibility loci, however, the mechanisms are unknown. We examined splicing QTLs (sQTLs) in islets from 399 donors and observed that genetic variation has a widespread influence on splicing of genes with important functions in islet biology. In parallel, we profiled expression QTLs, and used transcriptome-wide association and co-localization studies to assign islet sQTLs or eQTLs to T2D susceptibility signals that lacked candidate effector genes. We found novel T2D associations, including an sQTL that creates a nonsense isoform in *ERO1B*, a regulator of ER-stress and proinsulin biosynthesis. The expanded list of T2D risk effectors revealed overrepresented pathways, including regulators of G-protein-mediated cAMP production. This data exposes an underappreciated layer of genetic regulation in pancreatic islets, and nominates molecular mediators of T2D susceptibility.

## Introduction

Genome-wide association studies have identified hundreds of genomic loci that carry genetic variants contributing to Type 2 Diabetes (T2D) susceptibility^1–3^. The vast majority of associated genetic variants are non-coding, and epigenomic studies have revealed that many are located in human pancreatic islet transcriptional cis-regulatory elements^4–7^. Numerous T2D risk loci have thus been assigned to effector transcripts through human islet expression quantitative trait loci (eQTLs), three-dimensional chromatin maps, and genome editing experiments^8–11^. Such studies have established that gene expression variation in pancreatic islets is critically important for T2D susceptibility. A large fraction of T2D risk loci, however, cannot be ascribed to transcriptional regulatory mechanisms in pancreatic islets or other tissues, pointing to additional poorly understood genetic mechanisms.

Alternative splicing of pre-mRNAs provides an additional mechanism whereby genetic variation can create functional diversity across human genomes. Rare mutations and common variants that influence pre-mRNA splicing have been linked to a broad range of human diseases^12–14^. The effects of genetic variants on pre-mRNA splicing in human pancreatic islets, however, are largely unexplored. GTEx, the most comprehensive catalogue of splicing QTLs (sQTLs) across human tissues, does not include human pancreatic islets^14^. Other studies have examined exon-level expression in islet RNAs^10,15,16^, although this does not capture the complexity of alternative splicing and is confounded by total gene expression variation. We have now directly examined RNA splicing in a panel of 399 human islet samples and created an atlas of sQTLs. This showed that sQTLs impact key genes for islet biology and diabetes. We uncovered sQTLs that show fine colocalization with genetic variants associated with T2D, discover new genetic associations, and expand the spectrum of putative gene effectors of disease susceptibility. These findings provide fresh biological insights into genetic mechanisms underlying diabetes risk.

## Results

To examine the impact of common genetic variation on RNA splicing in pancreatic islets, we aggregated RNA-seq and genotype data from 447 human pancreatic islets samples from four cohorts^9,11,15,16^ and processed them to yield 399 qualifying samples after applying genotype and RNA-seq quality controls (**Figure 1a**). We corrected for known and unknown covariates (**Supplementary Figure 1**) and performed QTL analysis on mRNA levels (eQTLs) and junction usage^17^ (sQTLs), using 6.46 million common variants. Focusing on genes expressed in >10% of samples in each cohort, we found *cis*-eQTLs in 3,433 genes (eGenes) at FDR ≤ 1% (**Figure 1a**, **Supplementary Table 1**). Islet-expressed genes also revealed a widespread impact of genetic variants on splicing variation, with 4,858 *cis*-sQTLs at FDR ≤ 1%, 25% of which showed >10% shift in splice site usage in reference vs. alternate alleles (**Figure 1a, b**, **Supplementary Table 2**). The 4,858 sQTL junctions included alternative usage of 5’ exons, 3’ exons, mutually exclusive or skipped exons, or influenced combinations of such splice variants (**Figure 1c-i**). The junctions mapped to 2,088 distinct genes (sGenes), of which ∼90% were known protein-coding genes, and ∼7% were lncRNAs (**Supplementary Figure 2a**).

**Figure 1.**
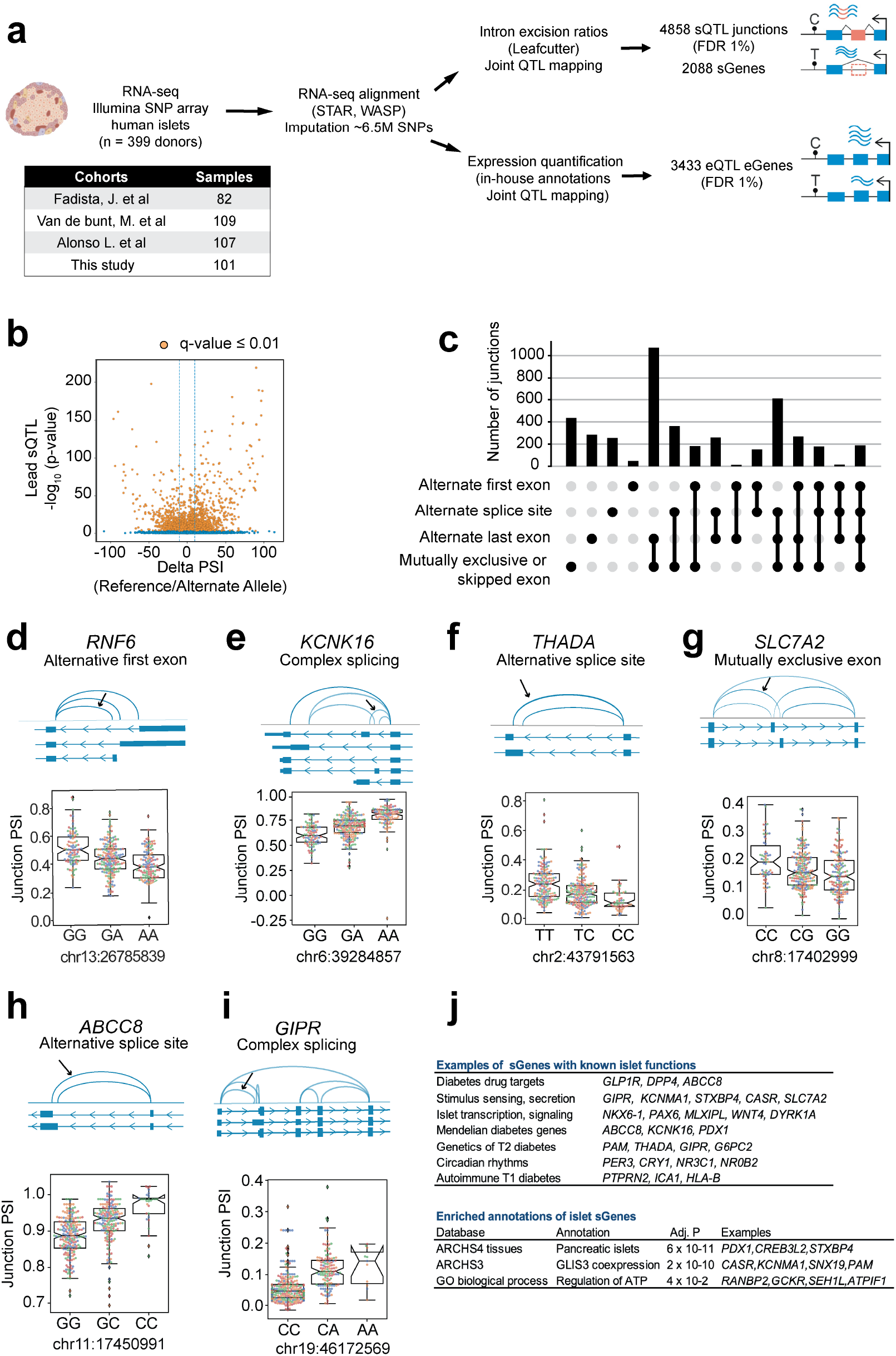
Mapping sQTLs and eQTLs in human pancreatic islets. **(a)** Overview of the study design. **(b)** Volcano plot showing the reference to alternate allele change in percentage splice index (Delta-PSI) for junctions, and sQTLs −log_10_ p-values. Orange dots depict sQTLs junctions with q ≤ 0.01. **(c)** Classification of sQTLs according to types of splicing events. **(d-i)** Selected examples of sGenes with different types of splicing events. An arrow signals the sQTL junction with best p-value, and adjacent boxplots show normalized, batch-corrected junction PSI values stratified by the lead sQTL genotype (IQR and 1.5 × IQR whiskers). Junction PSI values are colored according to the human islet dataset they belong to (see panel **Figure 1a). (j)** Functional annotations of sGenes. The top panel shows examples with known functions in islet function and diabetes, the bottom panel shows enriched annotations using EnrichR and Benjamini-Hochberg-adjusted p-values.

We benchmarked human islet splice variants against GENCODE^18^ and other available transcriptome maps and found that only 77% of the sQTL junctions were annotated. This overlap increased to 90% in comparisons with unpublished human islet transcript annotations built with long-reads (GA, AB, unpublished) (**Supplementary Figure 2b**). This suggests that human islet transcripts are still incompletely annotated, but nevertheless well captured by our analysis of splice junctions.

We compared islet sQTLs to previously reported exon-QTLs^10^, a possible proxy for sQTLs, and found that only 18% of sQTL junctions were flanked by exons from exon-QTLs. Furthermore, when sQTL and exon-QTLs affected the same gene, there was limited linkage disequilibrium (r^2^ < 0.6) between the lead sQTL and exon-QTL variant for 45.2% of overlapping genes (**Supplementary Figure 2c, d**). This indicates that sQTLs, which directly measure splice junction variation, and exon-QTLs, which measure exon levels and can thus be influenced by variables unrelated to RNA splicing, capture fundamentally different events.

Islet sGenes were enriched in islet-specific co-expression networks (**Figure 1j**) and included numerous genes with well-established roles in islet biology and diabetes, including major drug targets (*GLP1R, DPP4, ABCC8*), regulators of hormone secretion (*SLC7A2, CASR, GIPR*), transcription (*NKX6-1, PAX6, MLXIPL/ChREBP*), signaling (*DYRK1A, WNT4*), or circadian rhythms (*PER3, CRY1, NR3C1, NR0B2*). Importantly, sQTLs also affected genes that harbor variants that cause monogenic diabetes (*PCBD1*, *KCNK16, ABCC8, PDX1*) or influence T2D risk (*THADA, PAM*), as well as genes that play a role in T1D pathogenesis (*ICA1, PTPRN2, HLA-B*) (**Figure 1d-j**, **Supplementary Figure 3, Supplementary Table 2**). Our results, therefore, disclosed a pervasive impact of common genetic variants on alternative splicing of human pancreatic islet transcripts, including numerous genes that are important for islet function and differentiation, diabetes treatment, or pathophysiology.

### sQTLs and eQTLs are distinct

We next explored the extent to which genetic effects on splicing and expression of islet transcripts were distinct. Only 34% of genes that harbored sQTLs (715 sGenes) also harbored a significant eQTL (**Figure 2a**). Furthermore, for those 715 common genes, the lead eQTL and sQTL were frequently not in linkage disequilibrium (r^2^ <0.6 for 57% of genes, <0.1 for 24% of genes, **Figure 2b**). Thus, in most genes that harbored both eQTL and sQTLs these were driven by independent signals. This is illustrated by *RGS1*, encoding a regulator of G protein signaling, which has independent variants affecting either mRNA expression or exon skipping (**Figure 2c**). In keeping with these findings, eQTLs and sQTLs were enriched in different functional genomic annotations. sQTLs were predominantly enriched in 5’ or 3’ splice sites and exons, whereas eQTLs showed a predominant enrichment in active promoters and enhancers (**Figure 2d, Supplementary Table 3**). We also found differences in the extent to which genetic effects on splicing or expression differed across tissues; ∼60% of lead islet sQTLs showed significant sQTLs in <5 GTEx tissues^14^, compared to ∼30% of lead eQTLs, suggesting a significant islet-specific component of sQTLs (**Figure 2e**). Taken together, our results reveal two separable layers of genetic regulation of the human islet transcriptome.

**Figure 2.**
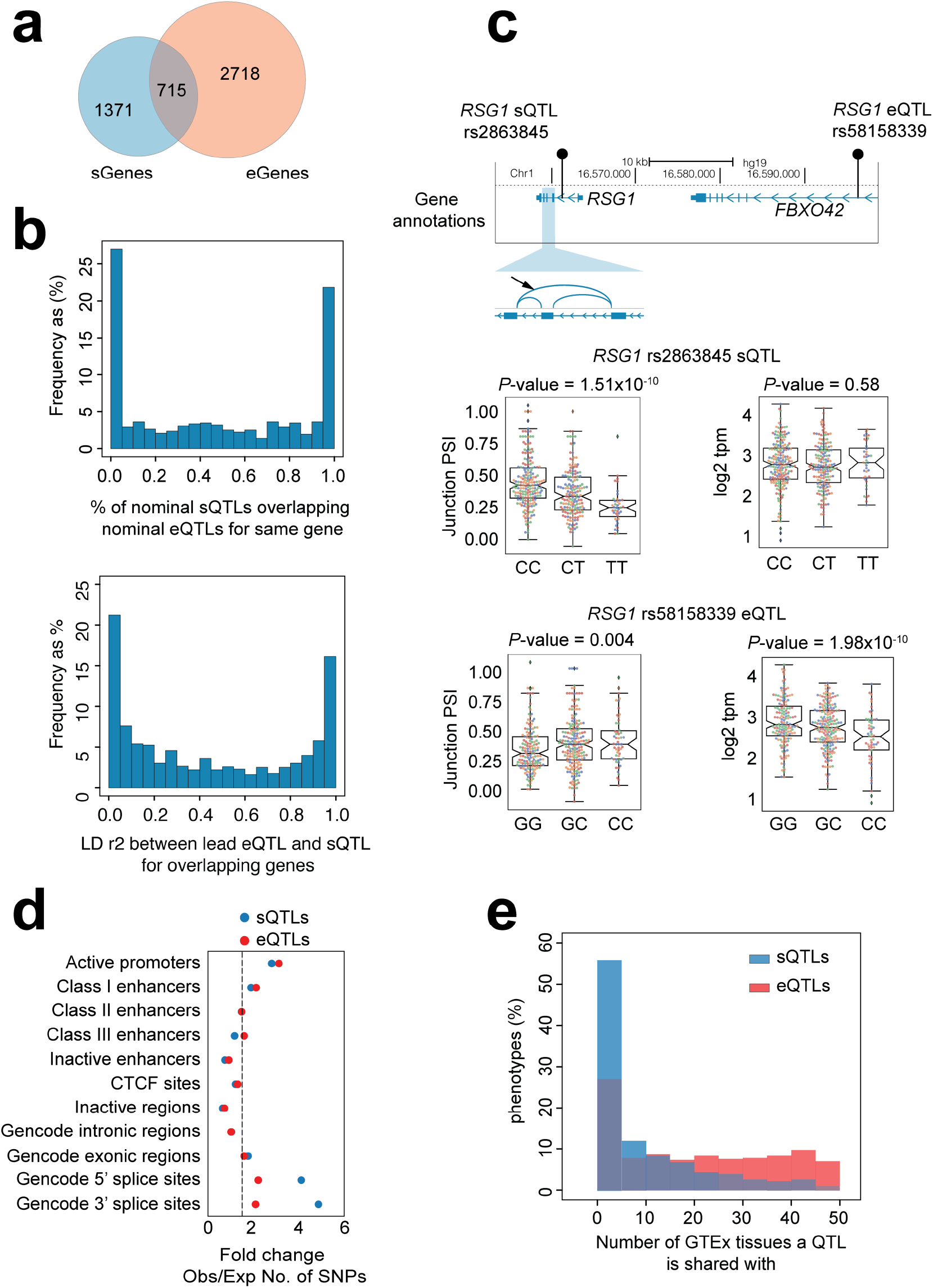
sQTLs and eQTLs are distinct genetic signals. **(a)** Overlap of sGenes and eGenes. **(b)** For 715 genes that have both eQTLs and sQTLs (overlapping genes in **Figure 2a)** the top histogram shows the distribution of the percentage of variants shared between sets of nominally significant eQTLs and sQTLs. The bottom histogram shows the distribution of LO (r^2^) values between the lead eQTL and sQTL. **(c)** *RSG1* has a distal eQTL, located in an intron of the *FBX042* gene, and an intronic sQTL, both of which are in low LO (r^2^=0.25). Boxplots represent *RSG1* expression and junction PSI values for both sQTL and eQTL, showing that the lead eQTL rs58158339 is not associated with *RSG1* splicing and the sQTL rs2863845 is not associated with expression. Boxplots show normalized, batch-corrected expression or junction PSI values stratified by the genotype of the lead QTL (IQR boxes and 1.5 × IQR whiskers). Individual samples are colored according to the human islet dataset they belong to. Nominal QTL p-values are provided. **(d)** Enrichment of sQTL and eQTL variants in different functional genomic annotations. The x-axis represents GREGOR fold change of observed vs. expected number of SNPs at each functional annotation. The dotted line represents 1.5-fold change. **(e)** Percentage of eGenes and junctions with e and sQTLs, respectively, shared in different number of tissues from the GTEx VS release.

### Islet sQTLs provide new T2D and glycemic trait targets

Genetic susceptibility for T2D has been linked to sequence variants that influence gene transcription in human islets^8–11^, but the relationship with islet splicing has not been systematically explored. To examine the potential contribution of islet sQTLs to T2D genetic associations^1^, we first used quantile-quantile plots that compare the distribution of T2D association p-values of sQTL and eQTL variants against an expected null distribution (**Figure 3a**). As anticipated, eQTLs showed a strong inflation of more significant T2D association p-values. Remarkably, sQTLs also showed genomic inflation of T2D risk p-values (**Figure 3a**). Importantly, this effect was maintained with islet-selective sQTLs (junctions with sQTLs in ≤ 5 GTEx tissues), or after omission of sQTLs that had high linkage disequilibrium with eQTLs (lead sQTL is in r^2^ ≥ 0.6 with the lead eQTL for the same gene) (**Supplementary Figure 4a,b**). Furthermore, sQTLs also showed genomic inflation of low association p-values with T2D-related traits such as fasting glycemia (FG) or fasting insulin (FI)^19^ (**Supplementary Figure 4c,d**). These observations, therefore, suggest that splicing variation in human islets could drive a component of T2D genetic susceptibility.

**Figure 3.**
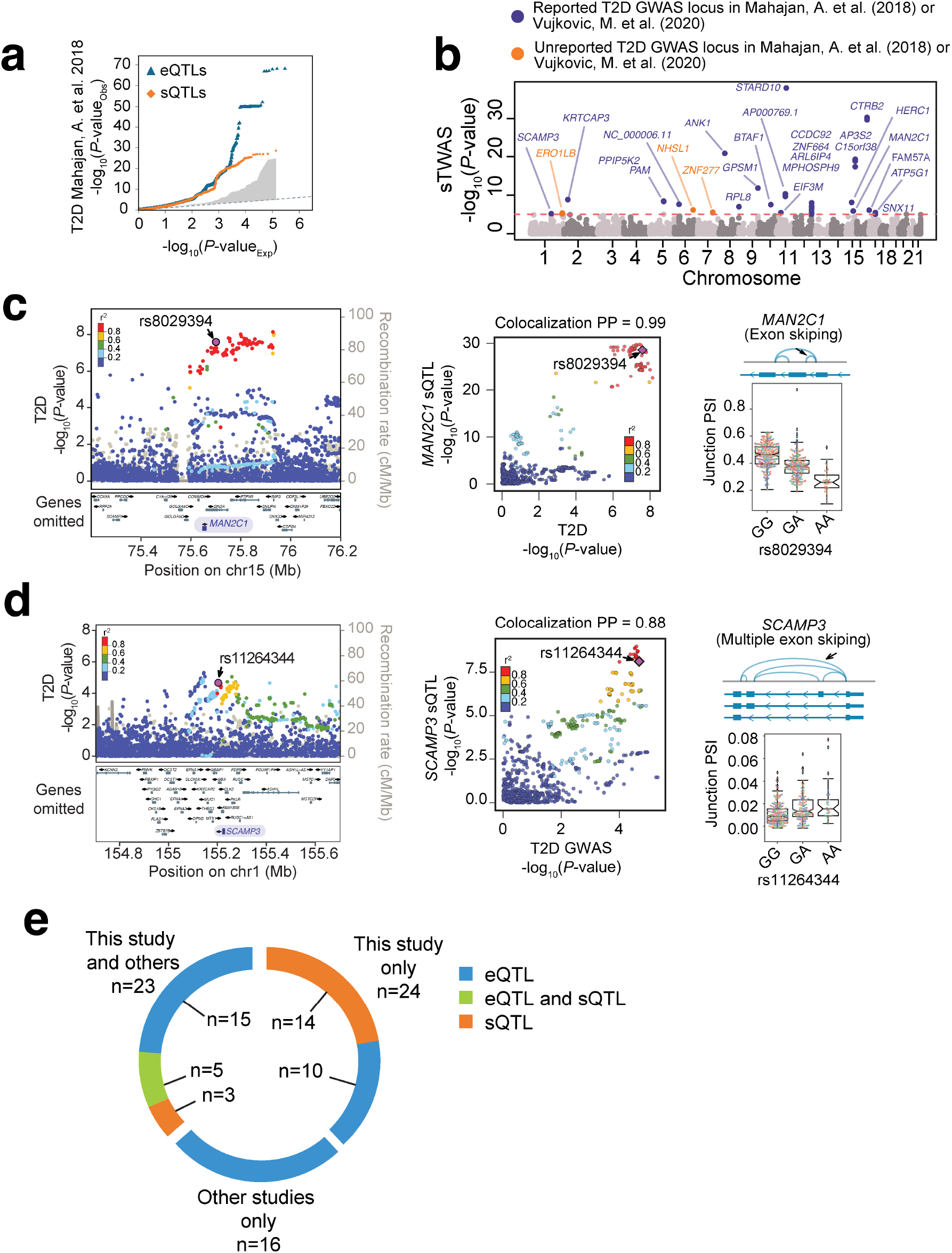
Role of human islet splicing variation in T2D susceptibility. **(a)** Quantile-quantile (QQ) plot showing observed T2D association p-values in human islet sQTLs (orange dots) and eQTLs (blue dots) against p-values under the null hypothesis. The grey shaded region represents observed −log_10_ p-values across 1000 random samplings of control sets of variants, with the same size as islet sQTLs. **(b)** Manhattan plot of splicing associations with T2D susceptibility (sTWAS). The y-axis shows −log_10_ TWAS association p-values. Significant sTWAS associations in known T2D GWAS loci are colored in purple, and in previously unreported loci in orange. **(c,d)** Regional T2D GWAS signal plots for *PTPN9* and *PKLR* loci, two known T2D susceptibility regions, which showed significant sTWAS signals in *MAN2C1* **(c)** and *SCAMP3* **(d)** genes, respectively. LocusZoom plots show −log_10_ T2D association p-values and locations in hg19 genome build. LocusCompare scatter plots show sQTL and T2D GWAS association p-values (−log_10_ scale), illustrating co-localization of variants for both traits. Variants are colored according to the LD correlation (r^2^) with the lead GWAS variant of the sTWAS association (purple diamond). Boxplots show normalized, batch-corrected junction PSI values on y-axis stratified by the genotype of the lead GWAS variant from the sTWAS association. **(e)** Known T2D loci with target effector transcripts nominated by sQTLs and eQTLs from this study and/or eQTL maps from the insPIRE consortium.

We reasoned that if splicing variation is truly instrumental in disease susceptibility at specific loci, a fraction of genetic signals for sQTLs and T2D association should show robust colocalization, and this could in turn point to specific transcripts underlying disease pathophysiology. To this end, we performed a systematic colocalization analysis between our islet sQTL or eQTLs, and independent GWAS signals for T2D (n = 403)^1^ or glycemic traits (fasting glycemia, FG/ fasting insulin, FI) (n = 274)^19^. We applied colocalization as implemented in *gwas*-*pw*^20^, which draws upon the original *coloc* algorithm but does not rely on user-defined priors. This identified candidate effector transcripts with robust evidence for colocalization (posterior probability of shared association between both phenotypes ≥ 0.8) at 9 independent T2D GWAS signals using sQTLs, and 25 using eQTLs (**Supplementary Tables 4, 5**). At loci associated with glycemic traits, we found robust colocalization with 8 putative target genes with sQTLs, and 17 genes with eQTLs (**Supplementary Tables 4, 5)**.

We further harnessed Transcriptome-Wide Association Studies (TWAS) to expand the collection of candidate effector genes for T2D and related traits. In this approach, which so far has not been applied to human islet RNA-seq data, genetic effects on splicing or expression are used to impute transcript variation in cases vs. controls from GWAS datasets. This allowed us to identify splicing or expression changes associated with T2D and related traits. More specifically, we used the FUSION algorithm^21^ and GWAS summary statistics^1^ to identify T2D associations with islet junction usage or gene expression (sTWAS and eTWAS, respectively). Because TWAS findings do not necessarily distinguish between shared genetic effects and linkage^22,23^, we focused on TWAS signals showing colocalization with the GWAS phenotype (PP4 ≥ 0.6), but minimizing confounding effects from linkage (PP3 < 0.8). This identified 27 genes (42 splicing events) showing significant sTWAS with T2D risk, and 29 genes with eTWAS after multiple testing correction (Bonferroni p= 8.6 × 10^−6^ and 1.8 × 10^−5^ after correcting for 5,804 splicing junctions and 2,851 genes, respectively) (**Figure 3b, Supplementary Figure 4e, Supplementary Figure 5, Supplementary Tables 6, 7**). For glycemic traits, we identified 22 candidate target genes (43 splicing junctions) of GWAS signals via sTWAS, and 16 candidate target genes via eTWAS (**Supplementary Figure 4f-i, Supplementary Tables 6,7**).

As expected, most TWAS signals fell in loci showing significant T2D and glycemic trait associations in GWAS **(Figure 3b, Supplementary Figure 4e-i, Supplementary Figure 6a-c, Supplementary Table 6,7)**. This included a sTWAS association for *MAN2C1*, encoding α-Mannosidase that has been implicated in mitochondrial-induced apoptosis and tissue damage^24,25^ (**Figure 3c)**. However, sTWAS revealed six T2D associations that did not reach genome-wide significance in the reference GWAS^1^, three of which (*SCAMP3, SNX11*, and *FAM57A*) were nevertheless significant in a recent trans-ancestral meta-analysis for T2D^3^ (**Supplementary Table 6**)*. SCAMP3* encodes a vesicular transport protein^26^ with unknown function in islets (**Figure 3d**). Other significant sTWAS for T2D did not reach significance in GWAS reported so far, namely those encoding *ERO1B, NHSL1* and *ZNF277* (**Figure 3b, Supplementary Table 6;** further details for *ERO1B* are shown in **Figure 4**). Thus, sTWAS nominated putative effector targets for T2D and glycemic trait genetic associations, and identified additional genetic associations.

Our studies also highlighted significant eTWAS for T2D, including *PCBD1*, which is mutated in a syndrome that includes monogenic diabetes^27,28^, and encodes a co-factor of HNF1A, another monogenic diabetes gene^29^ (**Supplementary Figure 6c, Supplementary Table 7**). This locus was not significant in the reference GWAS^1^ but another recent meta-analysis detected a significant association in this locus^3^. Our findings indicate that *PCBD1* is a strong candidate effector transcript for this T2D susceptibility locus. Previously unreported T2D genetic association signals loci were found through eTWAS for *CTC*-*228N24.2* genes and *VSNL1*, a Ca+ sensor that modulates cAMP and insulin secretion^30^ (**Supplementary Figure 6d, Supplementary Table 7)**.

Taken together, the combination of our TWAS and QTL colocalization results identified candidate effector transcripts for 47 known T2D susceptibility loci, including 22 acting through sQTLs, and 30 through eQTLs. Out of these 47 loci, 24 lacked information about likely effector transcripts in a recent large-scale islet eQTL analysis^10^. In aggregate, our sQTL and eQTL analysis increases the number of T2D loci with candidate effector genes supported by molecular QTLs to 63, a 1.6-fold change relative to the largest islet eQTL analysis so far^10^ (**Figure 3e, Supplementary Table 8**). Likewise, the integration of our molecular QTLs and GWAS summary statistics data of glycemic traits from a recent trans-ancestral meta-analysis^19^ allowed us to prioritize effector genes at 47 GWAS loci, a 3.8-fold increase relative to the largest islet eQTL mapping study so far (**Supplementary Table 9**).

### Fine-mapping QTLs identifies candidate causal variants for T2D risk

Genetic fine-mapping of molecular QTLs has been shown to not only highlight effector genes, but to also aid in the identification of causal variants for complex traits^31^. To this end, we generated 95% credible sets of sQTLs and eQTLs (**Supplementary Data 3,4**). For sQTLs, the highest credible set posterior probabilities (CPP) of fine-mapped variants were observed for markers in 5’and 3’ splices sites, followed by exonic and intronic regions (Mann-Whitney p = 1.4 × 10^−19^, 1.1 × 10^−21^, 4.5 × 10^−80^ and 1 × 10^−3^, respectively, compared with intergenic variants) (**Figure 4a**). Seqweaver, which provides variant effect predictions using RBP-based deep learning models^32^ showed enriched disease impact scores (DIS) of sQTLs in analogous annotations (**Supplementary Figure 7a**). Furthermore, fine-mapped intronic and exonic sQTLs disrupted motifs of auxiliary splicing regulators (*SRSF3, SRSF9*, *HNRNPA1, HNRNPC)*, while those near 3’ splice sites showed recurrent disruption of branch point motifs and core splicing components^33,34^ (**Supplementary Figure 7c**).

**Figure 4.**
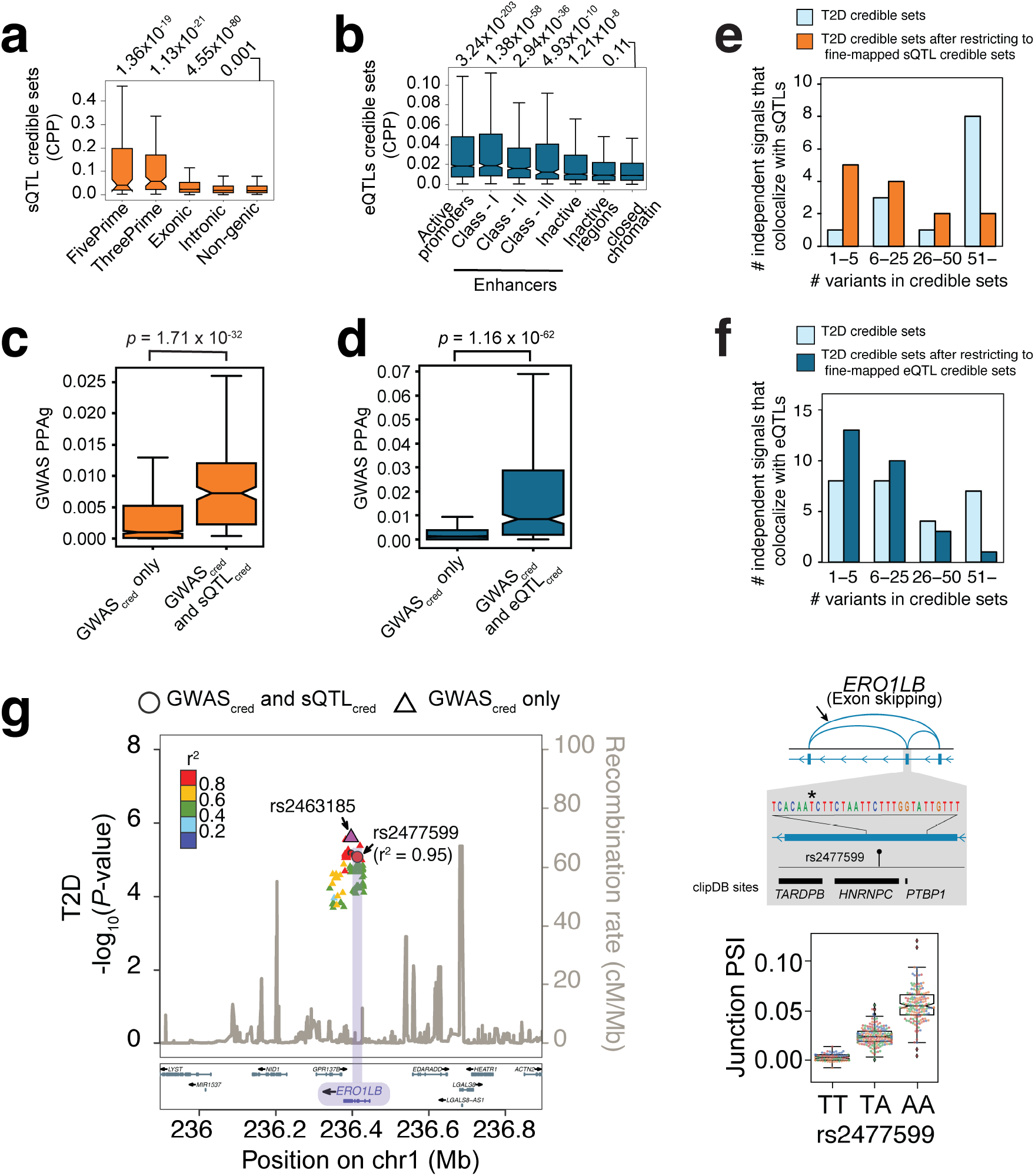
Fine-mapping causal variants for known and novel T2D genetic associations. **(a)** Distribution of sQTL causal posterior probabilities (CPP) across different genic and non-genic regions. P-values on top correspond to Mann Whitney comparisons with non-genic regions. **(b)** eQTL causal posterior probabilities across epigenomic annotations. P-values on top correspond to comparisons with credible set variants that fall outside islet epigenomic annotations (closed chomatin regions). **(c,d)** For all T2D-associated loci that colocalize with an islet QTL, we examined all fine-mapped variants (99% credible sets in GWAS, GWAS_cred_), and compared the distribution of T2D causal posterior probabilities for variants that are also fine-mapped QTL variants (QTL_cred_) vs. those that were not fine-mapped QTL variants. Mann-Whitney p-values are provided. Boxplots show IQR without outliers although p-values were calculated using all data points. **(e,f)** Integration of T2D GWAS credible set variants with credible sets from colocalizing sQTL and eQTL 99% credible sets increases fine mapping resolution. Bar plots show the number of independent signals that fall into different bins of number of candidate causal variants before and after restricting for QTL variants. **(g)** Fine-mapping an sQTL and T2D association in *ER01B*. The LocusZoom shows T2D association −10g_10_ p-values, credible set variants for GWAS and sQTLs are shown as circles, and other GWAS credible set variants as triangles. The color of dots reflects r^2^ with the lead GWAS variant (in purple), and includes the best fine-mapped sQTL candidate causal (rs2477599). The bottom inset depicts the alternative splicing event, along with the candidate causal sQTL variant and clipDB RBP binding sites. Boxplots are as described in **Figures 1–3**.

These orthogonal analyses were consistent with known determinants of RNA splicing, and highlight the potential of our fine-mapped sQTLs to prioritize causal variants.

Analogously, genetic fine-mapping of eQTLs revealed higher CPPs for variants in islet active promoters and enhancers (Mann-Whitney p= 3.24×10^−203^, 1.38×10^−58^, in promoters and Mediator-enriched enhancers, respectively) (**Figure 4b**). Similar enrichments were obtained with DeepSea^35^ (Supplementary Figure **7b**), and for disruption islet TF motifs^36–38^(**Supplementary Figure 7d)**. These results again supported that fine-mapped QTL variants have increased likelihood of driving splicing and expression variation in human islets.

Next, we hypothesized that if fine-mapped QTL variants are truly enriched in causal T2D variants, they should converge with variants that have highest CPPs in GWAS fine-mapping studies. To investigate this, we examined 99% credible set variants from T2D GWAS signals^1^ with colocalizing QTLs (PP4>0.8, 16 loci for splicing, 28 loci for expression). We observed that GWAS credible set variants that were also in sQTL credible sets had higher GWAS CPP than GWAS credible variants that were not in the sQTL credible sets (Mann-Whitney p= 1.71 × 10^−32^) (**Figure 4c**). Likewise, GWAS credible set variants that were also in eQTL credible sets showed higher GWAS CPP than other GWAS credible set variants (Mann-Whitney p= 1.16 × 10^−62^) (**Figure 4d**). This convergence between GWAS and QTL fine-mapping provided further evidence that QTLs truly contain causal T2D risk variants. The integration of QTL and GWAS credible sets increased the number of associated loci with ≤5 putative causal variants from 1 to 5 loci by integrating fine-mapped sQTLs, and 8 to 13 loci with fine-mapped eQTLs (**Figure 4e, f, see also examples in Figure 4g, Supplementary Figure 7e**). Thus, fine mapping QTLs has potential to boost the genetic resolution of GWAS credible sets.

*ERO1B* represents a salient example in which we fine-mapped a putative causal variant for a candidate effector. *ERO1B* showed a significant sTWAS T2D association (p= 6.5 × 10^−6^) in our studies, and only suggestive associations in GWAS^1^ (p= 2.3 × 10^−6^). Fine-mapping highlighted an exonic rs2477599 variant that localizes to a splicing silencer motif *HNRNPC*, which causes an exon skipping event that results in premature truncation of the *ERO1B* open reading frame (**Figure 4g)**. Previous genetic loss-of-function studies have shown that *ERO1B* (endoplasmic reticulum oxidoreductase 1 beta, also known as *ERO1LB*) is an ER protein that plays a critical role in insulin biosynthesis and β-cell survival^39–41^. This locus has not been previously reported in T2D GWAS studies, but FinnGen biobank data shows a significant T2D association with an intronic variant (rs1254190; p = 9 × 10^−9^, data freeze 5) that is only moderately linked to the sQTL (r^2^ = 0.50), suggesting that different alleles could alter T2D risk through *ERO1B*. Collectively, these lines of evidence highlight a T2D susceptibility locus at *ERO1B*, with a fine-mapped putative causal splicing variant and a plausible effector mechanism for T2D susceptibility.

### Islet QTLs provide insights into T2D pathways

The availability of an expanded list of candidate effector genes, as opposed to lists of genes that are located in the vicinity of associated SNPs, allowed us to explore the hypothesis that a subset of genetic signals could influence T2D susceptibility by acting on specific cellular pathways in pancreatic islet cells. We thus compiled 106 putative effector genes for T2D or glycemic traits (FG, FI) reported here as well as previously reported co-localizing islet eQTLs^10^. This list excluded those exclusively detected in non-endocrine cells after analysis of single cell RNA-seq datasets (**Supplementary Table 10**), as well as non-coding transcripts and genes without known function. Functional gene annotations revealed notable enriched pathways, including regulators of fatty acid biosynthesis (SCD5, FADS1, GCDH, BDH2, PAM; GO:0006636, GO:1901570, ENRICHR adjusted p = 0.04), and genes upregulated by hypoxia or mTORC1 activation (MsigDB Hallmark Hypoxia and mTORC1 signaling; q < 0.05 and < 0.001, respectively). We further used the list of 106 genes to identify networks using STRING ^42^ v11.5, using co-expression, experimental and functional annotation databases and allowing for minimal network inflation (≤5 interactions), while omitting text-mining to preclude bias arising from publications that name genes at significant GWAS loci. The resulting network exhibited 1.8-fold greater protein-protein interactions than random sets (PPI enrichment p < 10^−4^) and contained two distinct subnetworks (**Figure 5a**). One subnetwork contained components of the eIF3 translational initiation complex (FDR = 0.018). Several monogenic diabetes genes target translational initiation, in particular eIF2 complex (*EIF2B1, EIF2S3, EIF2AK3)*, due to their impact on endoplasmic reticulum stress in β cells^43,44^ (**Figure 5a**). This process, interestingly, is also regulated by ERO1B^41^. Another conspicuous sub-network was formed by molecular mediators of G protein-mediated enhancement of cAMP, a major insulinotropic pathway^45,46^. Manual curation revealed 8 genes in this pathway, including *GLP1R*, *RGS17*, *PDE8B*, four members of the GNAI3 (G(i) subunit α3) protein-protein interaction complex (*GPSM1, RGS19, ADRA2A, ADCY5*), and *VSNL1*, a modulator of cAMP and insulin secretion^30^ (**Figure 5a, 5b)**. These findings, therefore, provide clues to understand pathways and mechanisms that underlie islet dysfunction in T2D.

**Figure 5.**
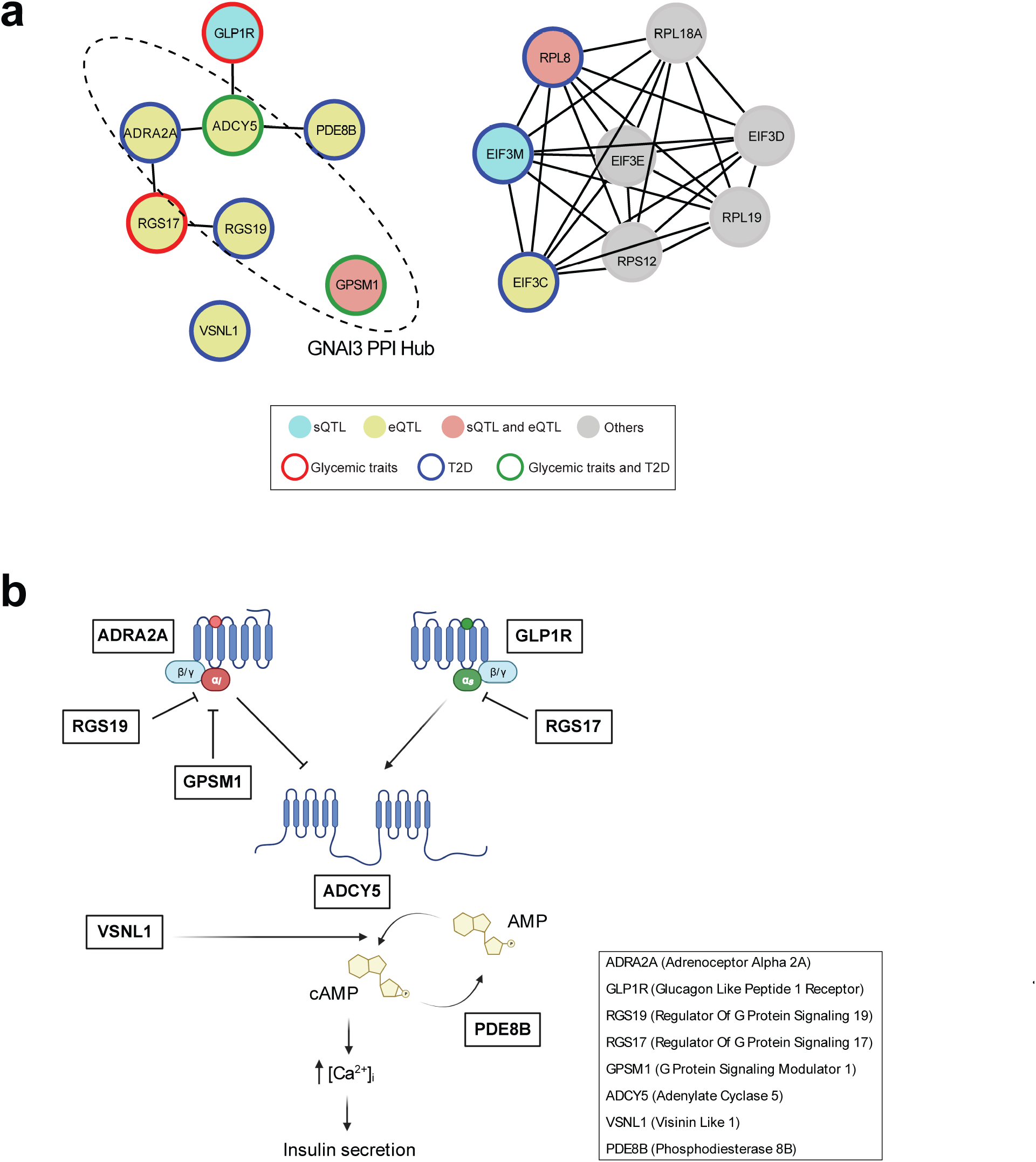
Co-localizing islet QTLs target distinct pathways. **(a)** STRING v11.5 was used to analyze 106 genes with sQTLs or eQTLs co-localizing with T2D or glycemic traits, including previously reported eQTLs. Only those with known or presumed protein-coding function, and not exclusively expressed in non-endocrine cells by single cell RNA-seq. We allowed for inflation of ≤ 5 interactors, and used default (>0.4) confidence scores. Shown are two networks with >3 components, one populated by components of heterotrimeric G protein signaling, and another by genes involved in elF3 translational initiation. Genes are colored as indicated in the label legend, other interactors added through STRING analysis are shaded gray. **(b)** Manual curation as used to illustrate the relationship between components of the network of G protein-mediated insulinotropic pathway genes targeted by islet QTLs linked to T2D and related traits.

## Discussion

This study adds splicing variation in pancreatic islets to the spectrum of molecular mechanisms that underlies T2D predisposition. Earlier studies had examined expression QTLs in human pancreatic islets^8–11,15,16^. The current study offers the first systematic analysis of how common genetic variants influence alternative RNA splicing in human islets. Parallel profiling of splicing and expression QTLs in the same dataset demonstrated that these represent two distinct mechanisms through which genetic variation can influence islet biology and disease. We found islet sQTLs that impact genes that have major roles in islet cell function as well as in the pathophysiology or treatment of various forms of diabetes. Furthermore, we observed a selective inflation of T2D association values among sQTLs, some of which showed fine co-localization with T2D variants. Finally, we applied for the first time splicing and expression TWAS to nominate T2D target genes, and identified novel T2D genetic association signals.

Gene targets that are nominated based on human genetic evidence double their likelihood of success in drug development pipelines^47^. We have expanded the current list of putative effector genes that mediate diabetes risk at known and novel susceptibility regions, and include examples that provide insights into T2D pathophysiology. This was illustrated by the association of T2D risk with a fine-mapped splice variant that creates a premature stop codon in *ERO1B*. This newly reported locus is supported by an independent T2D association signal observed in FinnGen, as well as prior experimental and genetic studies that point to a central role in ER homeostasis, insulin biosynthesis and diabetes progression^39–41^.

T2D has a highly polygenic architecture, with a very large number effector genes that individually exert small effects. It remains possible, however, that many such genes converge on a small number of biological pathways. The extended list of genes with islet QTLs that co-localize with GWAS signals allowed us to explore shared pathways underlying T2D susceptibility. We found supportive evidence for a role in translation, ER stress, and fatty acid metabolism, and, most clearly, 8 out of 106 QTLs that co-localized with glycemic trait or T2D associations were linked to G protein-coupled receptor (GPCR) signaling through cAMP. This pathway has been extensively involved in transducing signals from a vast range of extracellular insulinotropic stimuli, including incretin hormones, neurotransmitters, nutrients such as fatty acids, or extracellular matrix components^45,48,49^. Effectors from this pathway included an sQTL for *GLP1R*, a major drug target for T2D^50^, four members of the same protein interaction complex, and a novel T2D association for *VSNL1*, previously shown to stimulate cAMP production and insulin secretion^30^. These findings, therefore, suggest that abnormal production of cAMP plays a causal role in T2D susceptibility, plausibly through its impact on insulin secretion and islet gene transcription. This pathway is a known therapeutic target to stimulate insulin secretion. However, our findings raise the additional prospect that it is possible to target deranged GPCR signaling in precision medicine strategies that aim to correct causal defects, and thereby modify disease progression in susceptible individuals.

Our analysis of molecular QTLs entails some limitations. Splicing activity was estimated by junction usage, which does not directly inform isoform level regulation, and therefore provides a partial picture of how local splicing translates into gene function. Furthermore, the analysis of molecular QTLs was limited to islets, where genetic variants are known to play a central role in diabetes mellitus, most clearly in T2D. Some disease associations are expected to be mediated through other cell types and the mediating mechanism is not captured by our studies. Finally, colocalization and TWAS evidence does not preclude horizontal pleiotropy, and we have thus not unequivocally linked non-coding variant disease associations with the causal molecular targets. Instead, we have nominated plausible candidate effectors from a tissue with strong disease-relevance, which requires additional follow-up with orthogonal genetic and experimental studies to establish causality.

In addition to the discovery of causal genes and pathways for T2D risk, the splicing QTL resource reported here holds relevance for other diabetes-relevant contexts. For example, islet sQTLs can also be considered to understand genetic variants that modify autoimmune T1D risk. The genetic risk for T1D is largely mediated through effects on immune cells^51,52^, although recent studies have highlighted a role of genes expressed pancreatic exocrine and endocrine cells in T1D pathophysiology^53–56^. We identified examples of islet splicing QTLs at *MEG3* lncRNA and *ESYT1* that co-localize with T1D association signals^55^ (**Supplementary Figure 8a,b**) and thereby expose new candidate biological mediators for T1D.

This resource of islet sQTLs is also potentially relevant for efforts to dissect multi-allele interactions. There is growing evidence that disease-associated haplotypes can contain more than one functionally interacting causal variant^57^. Islet splicing variants can thus act in *cis* with other functional variants to influence disease susceptibility. For example, sQTLs could modify the penetrance of causal coding or cis-acting variants in both polygenic or Mendelian settings. Finally, we have shown that islet sQTLs can alter drug target genes, and can thus be examined to understand how genetic variation alters the response to therapies. Taken together, our findings provide resources and new avenues to understand the genetic underpinnings of diabetes mellitus.

## Supporting information

Overview of Supplementary Tables

## Acknowledgements

This research was supported by Ministerio de Ciencia e Innovación (BFU2014-54284-R, RTI2018-095666-B-I00), Medical Research Council (MR/L02036X/1), a Wellcome Trust Senior Investigator Award (WT101033), European Research Council Advanced Grant (789055), EU Horizon 2020 TDSystems (667191), ESPACE (874710) and Marie Sklodowska-Curie (643062, ZENCODE). S.B.G was supported by a Juan de la Cierva postdoctoral fellowship (MINECO; FJCI-2017-32090). M.I. was supported by a European Research Council consolidator award (101002275). Human islets for research were supported by the European Consortium for Islet Transplanation (ECIT; Juvenile Diabetes Research Foundation grant 2-RSC-2019-724-I-X). Work in CRG was supported by the CERCA Programme, Generalitat de Catalunya and Centro de Excelencia Severo Ochoa (SEV-2016-0571). We thank the Center of Genomic Regulation and NIHR Imperial BRC Genomics Units. A.L.G. is a Wellcome Senior Fellow in Basic Biomedical Science, and was supported by the Wellcome Trust (095101, 200837, 106130, 203141), the NIDDK (U01DK105535 and UM1 DK126185) and the Oxford NIHR Biomedical Research Centre.

## Contributions

G.A., S.B.G., J.F. conceived, coordinated the study, interpreted results and wrote the manuscript with input from remaining authors. A.B. analyzed transcript annotations, I.M. provided guidance on transcriptome analysis during initial phases, M.C.A provided insights into pathway analysis and M.I. on RNA splicing. A.L.G and L.G. provided datasets and phenotypes, T.B. F.P, L.M., J.K-C, M.S., P.M, L.P. procured human islets. J.G-H processed RNA and DNA samples, G.A., S.B.G. processed genome data, designed and performed all genetic and statistical analysis. All authors approved the final manuscript.

## Data availability

Raw RNA and genotypes will be made available from the European Genome–phenome Archive (EGA; https://ega-archive.org/).

Full sQTL and eQTL results as well as variant effect predictions will be made available at https://www.crg.eu/en/programmes-groups/ferrer-lab#datasets

### Conflicts of Interest

ALG’s spouse is an employee of Genentech and holds stock options in Roche.

## Notes

### Competing Interest Statement

The authors have declared no competing interest.

