## Supplementary Figures for "Genetic regulation of RNA splicing in human pancreatic islets"

### Supplementary Figure 1

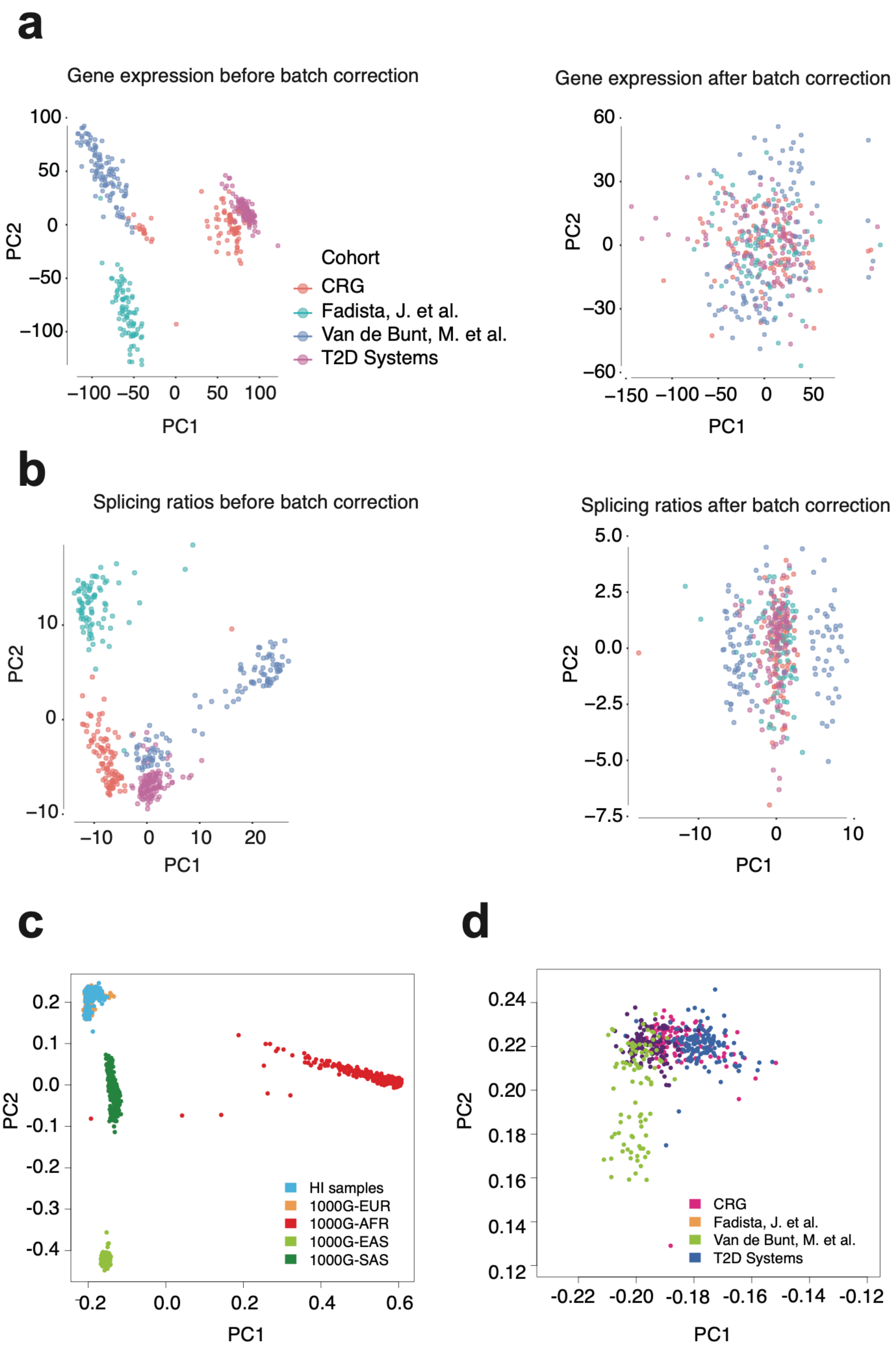

**Supplementary Figure 1. Principal component (PC) analysis based on RNA-seq and genotype data in 399 qualifying human islet samples. (a,b)** PCs calculated using gene expression and *Leafcutter* junction usage profiles before and after correcting for batch effects. **(c)** PCs calculated from genotypes. Islet sample donors (light blue) were positioned according to PCs from 1000 Genomes Phase 3 genotypes. **(d)** Differences in population structure between the four cohorts included in our panel of 399 islet transcriptomes.

Low resolution files for review

#### Supplementary Figure 2

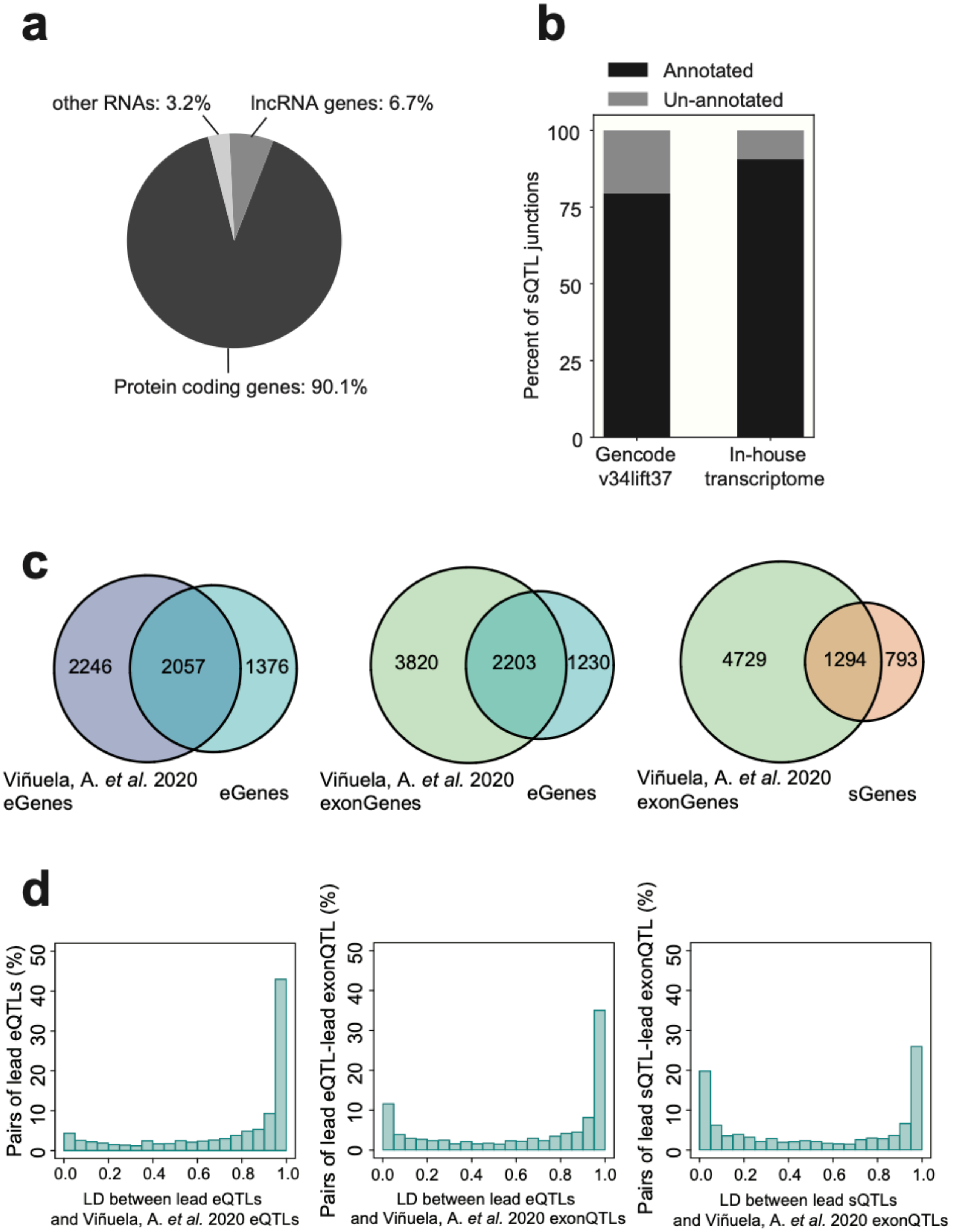

**Supplementary Figure 2. Characterization of sQTLs in human islets.**
**(a)** Distribution of sQTL junctions among protein-coding, lncRNAs and other RNAs.
**(b)** Percentage of sQTL junctions successfully annotated using GENCODE or unpublished in-house transcriptome annotations. **(c)** Venn-diagrams for the overlap between eGenes and sGenes identified in the current study vs. eGenes and exon-QTL genes identified by the InsPIRE consortium **(d)** Distribution of linkage disequilibrium ($r^2$) values between the lead eQTL or sQTL in our study and the lead eQTL or exonQTL in the insPIRE consortium in shared QTL genes**.**

### Supplementary Figure 3

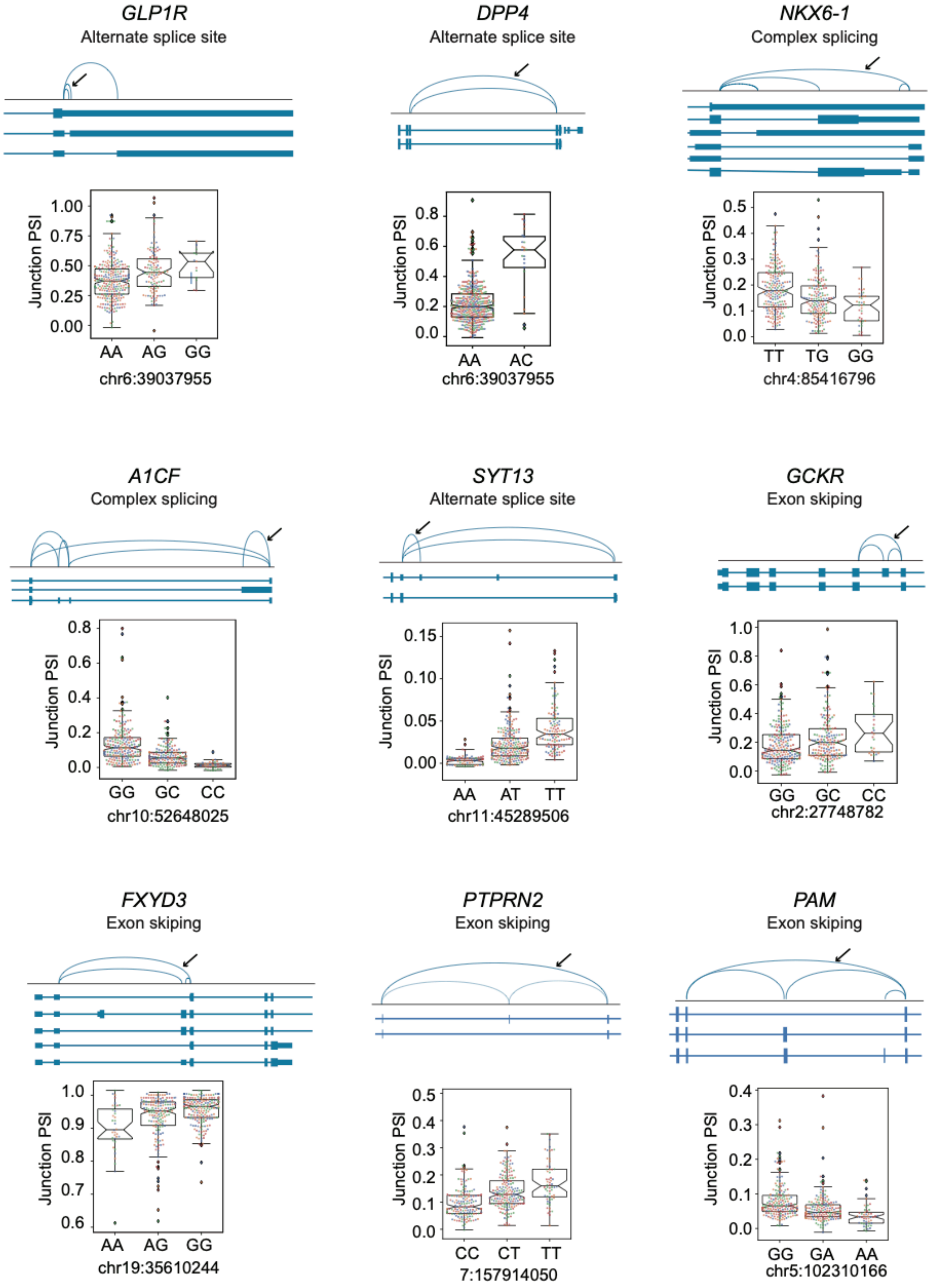

**Supplementary Figure 3. sQTLs at selected genes with central roles in islet cellular identity and diabetes pathophysiology**. Boxplot representations show the junction *percent spliced in* (PSI) values for the lead sQTL genotypes. The IQR of the junction PSI distribution is represented as boxes and whiskers are 1.5 x IQR. Individual PSI junction values are colored according to the human islet datasets used in this study.

### Supplementary Figure 4

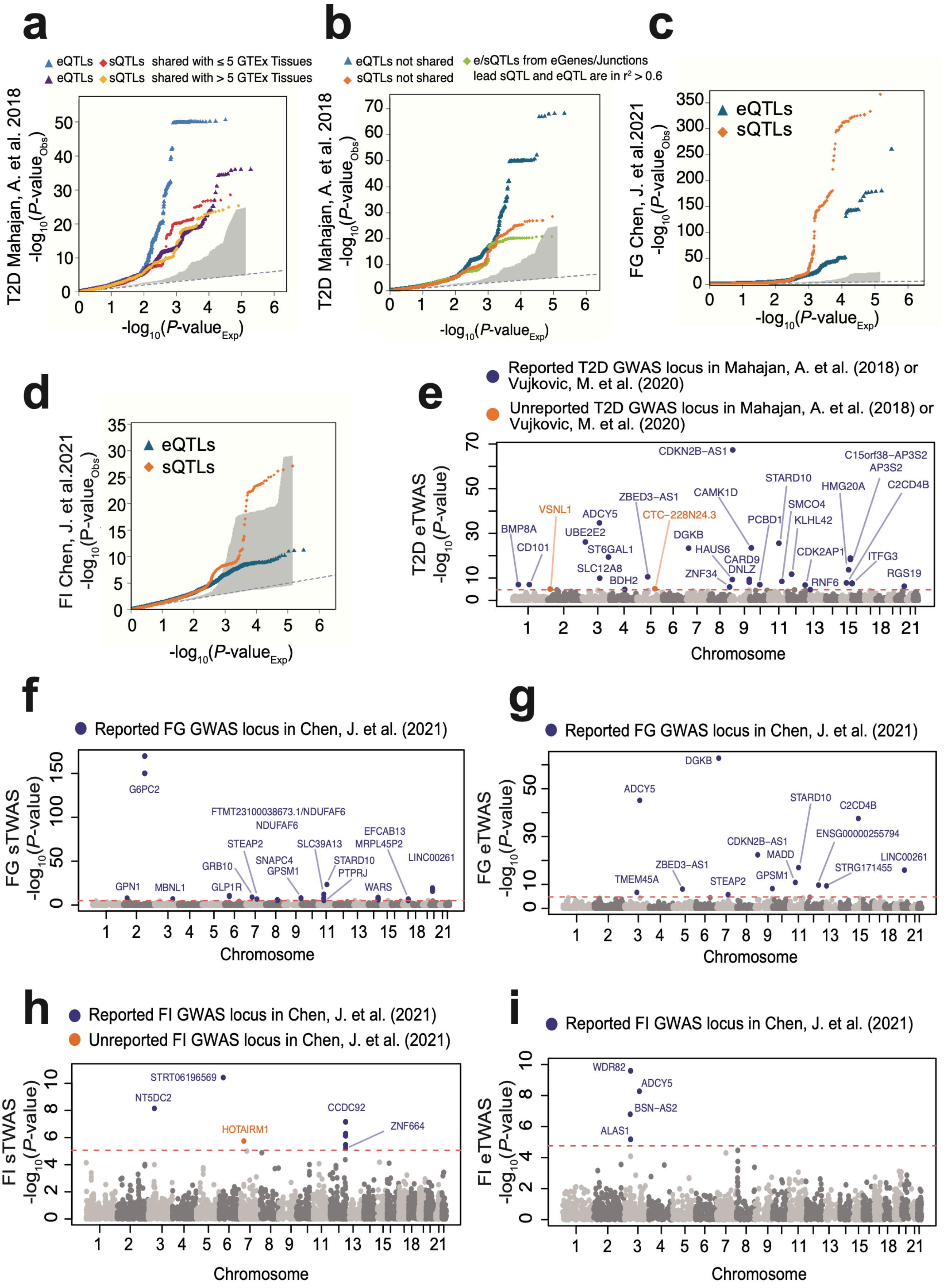

**Supplementary Figure 4. Role of islet sQTLs in T2D and related traits**.

**(a-d)** Quantile-quantile (QQ) plot representations show $-\log_{10}$ association p-values (y-axis) for sQTLs (orange) and eQTLs (blue), which are represented against the p-values under the null hypothesis (x-axis). The grey shaded area shows observed $-\log_{10}$ p-values across 1000 random samplings of control sets of variants, with the same size as islet sQTLs. **(a)** Inflation of T2D association p-values across eQTLs and sQTLs shared with either less or more than 5 GTEx tissues. **(b)** inflation of T2D association p-values for islet eQTLs and sQTLs that are independent vs. those in which a lead eQTL and sQTL are in $r^2 > 0.6$. **(c,d)** genomic inflation in fasting glycemia and fasting insulin association p-values for islet sQTLs and eQTLs. **(e-i)** Manhattan plots showing splicing (sTWAS) and gene expression (eTWAS) associations with T2D and variation of related traits. TWAS p-values are shown in the y-axis in the $-\log_{10}$ scale. Significant TWAS associations in known T2D GWAS loci are colored in purple while orange dots highlight significant TWAS associations in genomic regions that were not previously reported. FG = fasting glucose, FI = fasting insulin.

### Supplementary Figure 5

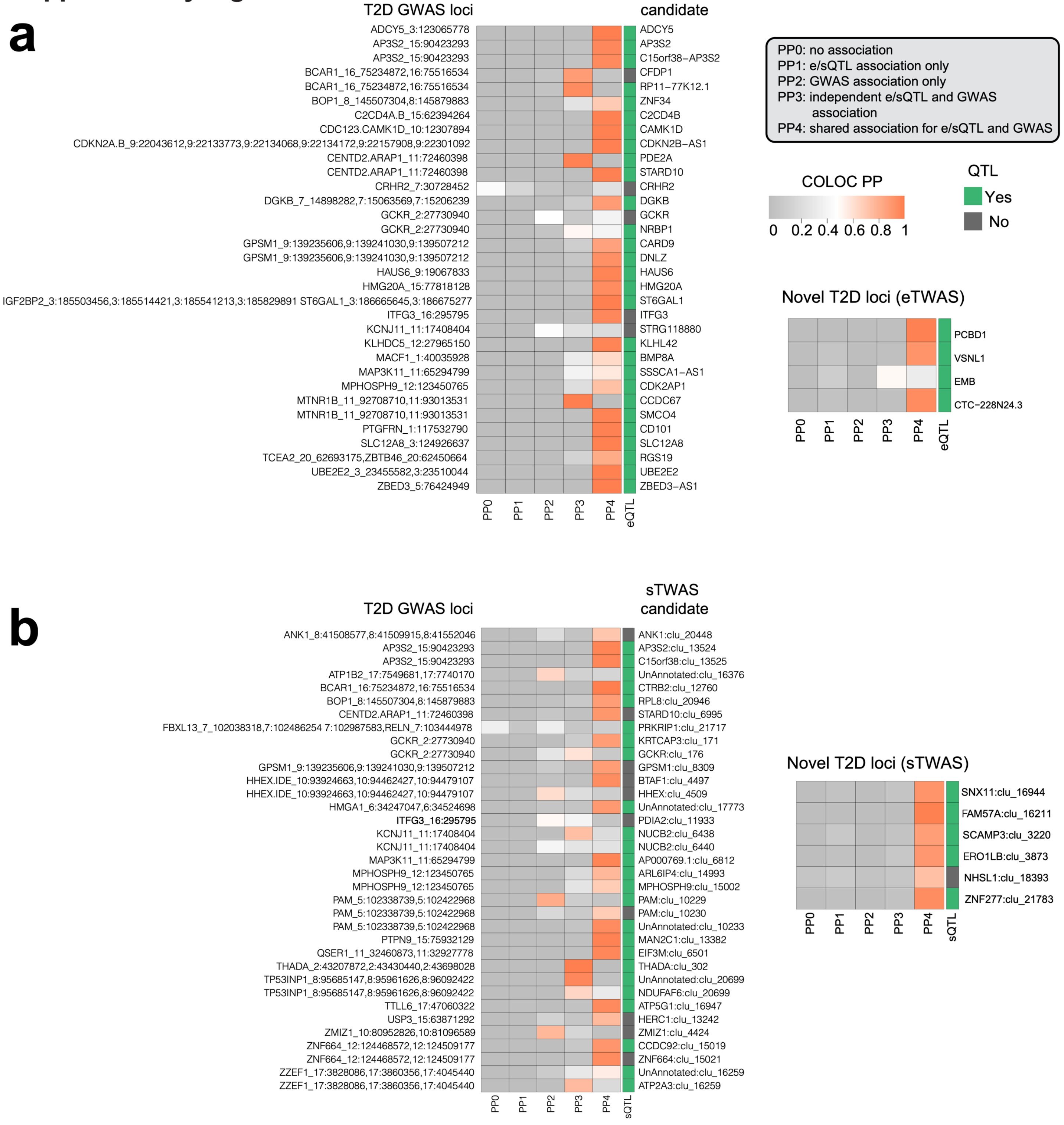

**Supplementary Figure 5. Colocalization posterior probabilities for TWAS associations. (a,b)** Heatmap representations show colocalization posterior probabilities from COLOC for **(a)** eTWAS and **(b)** sTWAS associations. Separate heatmaps are provided for known and novel T2D loci identified by TWAS. Heatmap color indicates five possible colocalization posterior probability values (PP0-PP4, see inset). TWAS-prioritized candidate genes that are also significant sQTL or eQTLs in human islets are highlighted in green.

Low resolution files for review

#### Supplementary Figure 6

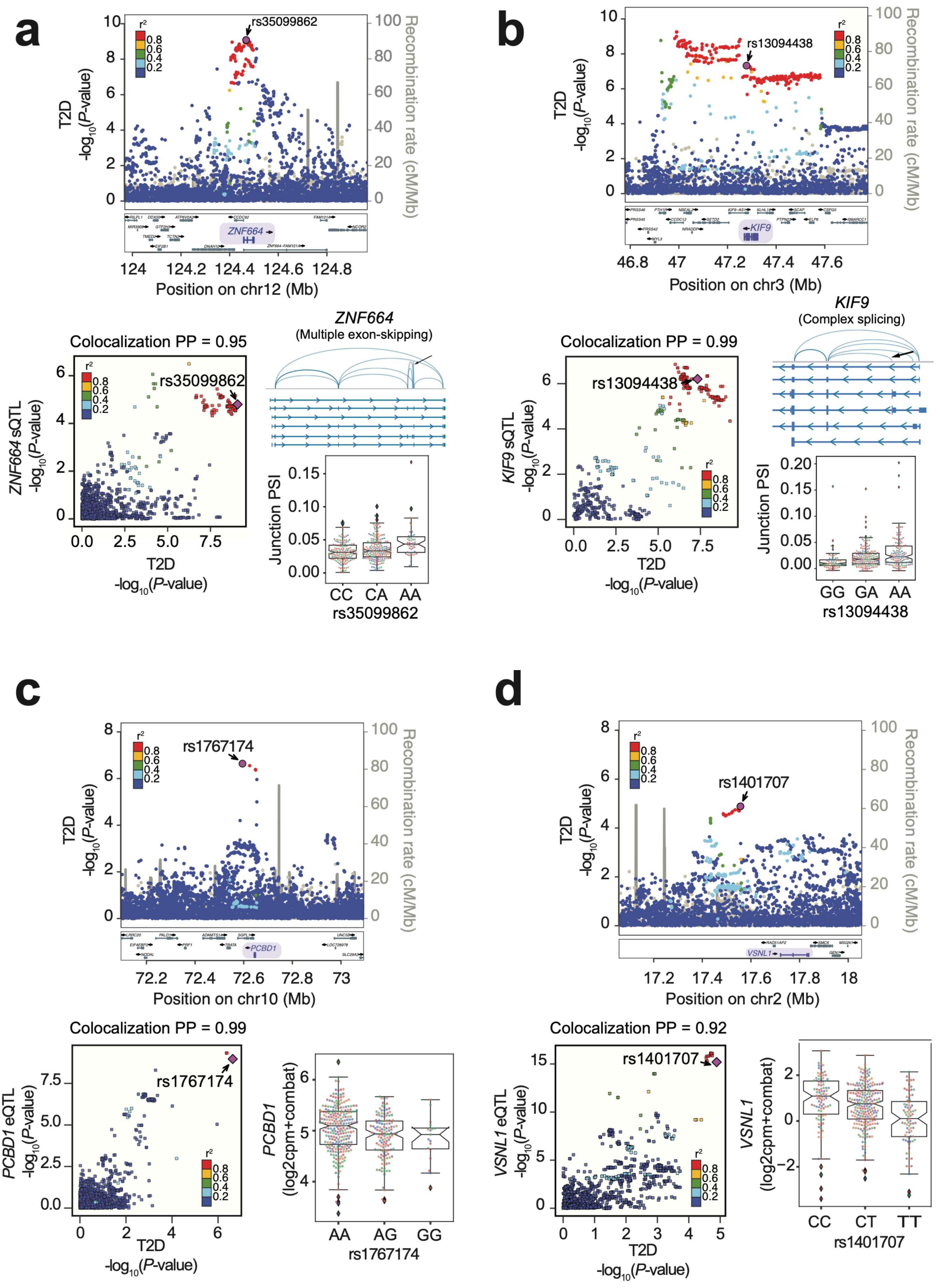

**Supplementary Figure 6. Regional signal plots for sTWAS and eTWAS associations mediating T2D risk**. **(a,b)** TWAS associations for alternative splicing in **(a)** *ZNF664* and **(b)** *KIF9* genes, which are located in known T2D susceptibility regions, as well as **(c,d)** eTWAS associations mediated by *PCBD1* expression in a known T2D locus, or for *VSNL1* in a previously unreported T2D association locus. For all panels, LocusZoom representations show -$\log_{10}$ T2D association p-values across the genomic position in hg19 genome build in the x-axis. Variants are colored according to the LD correlation ($r^2$) with the lead variant of the sTWAS association (purple). LocusCompare scatterplots show correlations between e/sQTL and T2D GWAS association p-values (-$\log_{10}$ scale), colored according LD correlations ($r^2$) with the lead GWAS variant from TWAS (purple) as the color scheme. Boxplots show PSI junction values or normalized gene expression values for the lead TWAS association variant genotypes.

#### Supplementary Figure 7

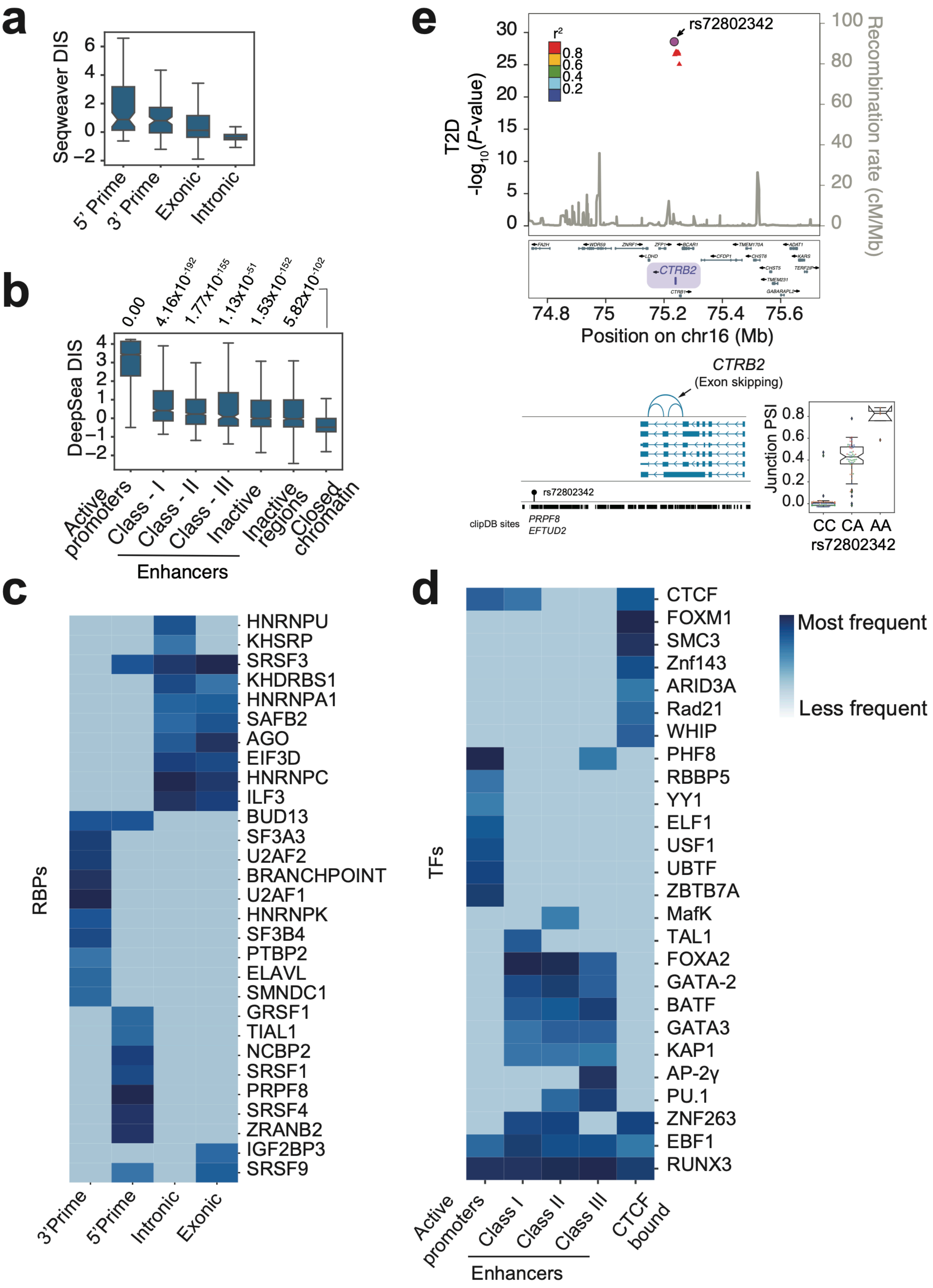

**Supplementary Figure 7. Functional annotation of fine-mapped sQTL and eQTL variants. (a)** Distribution disease impact scores (DIS) trained using Seqweaver in fine-mapped (credible set) sQTL variants across relevant functional annotations. **(b)** DIS trained using the DeepSea model in eQTL credible set variants located in human islet active promoters, enhancers, accessible chromatin regions lacking active modifications (*inactive* regions), and regions with *closed* (inaccessible) chromatin. **(c,d)** Disruption of DNA- and RNA- binding protein sequences by eQTLs and sQTLs. sQTL credible set variants show enriched disruption of largely expected sets of RBP sites according to their genomic location. Similarly, eQTL credible set variants show increased frequency of different TF binding sites according to their genomic location in different cis-regulatory elements. The frequency indicates the ranking of RBP/TF based on number of credible set variants impacting them with in each category. Top ranked RBP/TF with in each category are shown in dark blue. **(e)** Fine-mapping T2D associations and *CTRB2* sQTLs in the *BCAR1* T2D locus. Regional signal LocusZoom plot shows T2D association $-\log_{10}$ p-values and locations in the hg19 genome build, credible set variants for GWAS and sQTLs are represented as circles, and other GWAS credible set variants as triangles. Variants are colored according the $r^2$ with the lead and best candidate causal GWAS variant (in purple). The bottom inset depicts the alternative splicing event, along with the candidate causal sQTL variant and clipDB RBP binding sites where it is located. Boxplots are as described in **Supplementary Figure 6**.

### Supplementary Figure 8

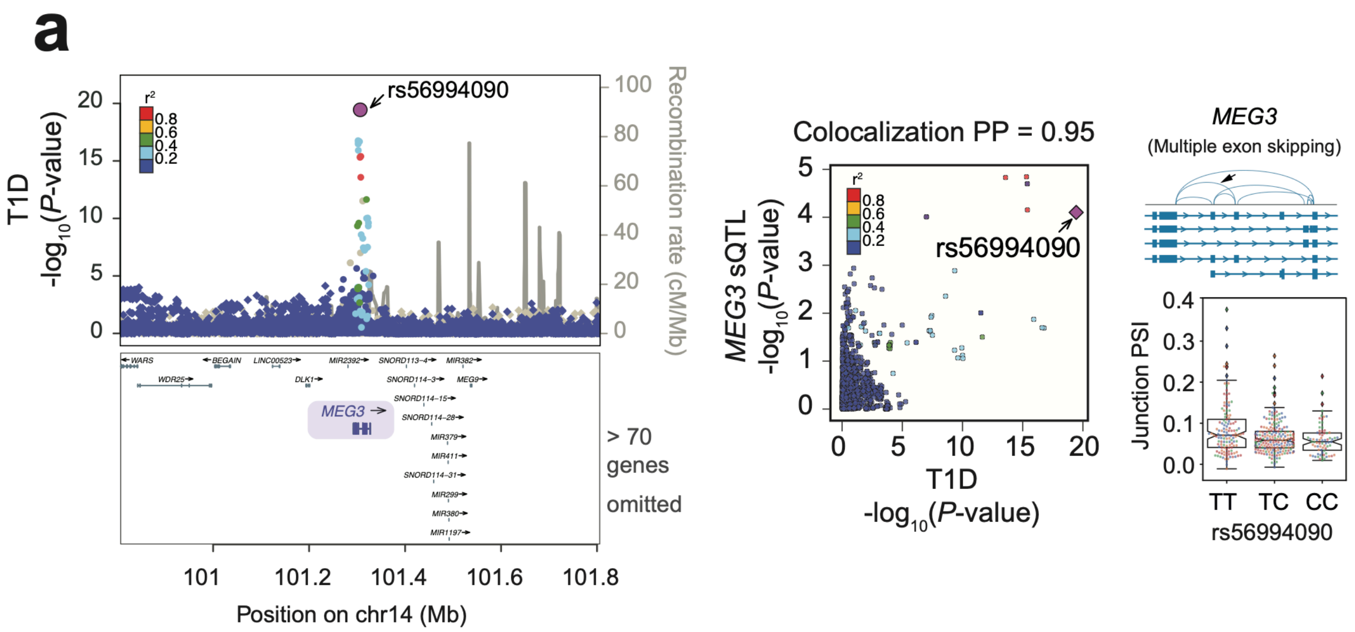


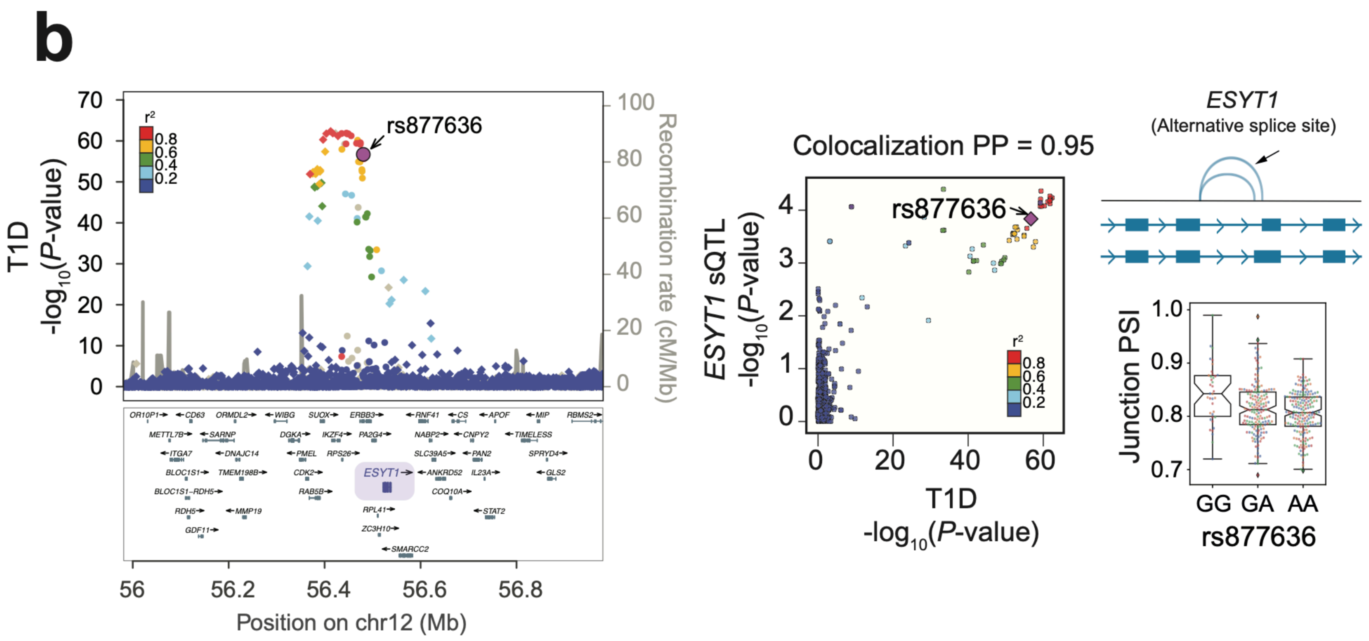

**Supplementary Figure 8. Co-localization of islet molecular QTLs with T1D predisposition signals**. **(a,b)** LocusZoom plots showing -$\log_{10}$ T1D association p-values at two known T1D loci, *DLK1/MEG3* and *IKZF4*, in which we identified colocalization with *MEG3* and *ESYT1* sQTLs using *gwas-pw* and *COLOC*, respectively. LocusCompare scatter plots illustrate correlations between sQTL and T1D GWAS association p-values (-$\log_{10}$ scale). Variants were colored following the linkage disequilibrium ($r^2$) with the lead GWAS variant (purple). Boxplots show PSI junction values according to individual sQTL genotypes for the lead GWAS association. GWAS summary statistics from a recent meta-analysis for T1D from Chiou, J. and colleagues (2021) have been used.
