## Supplementary material for "Genetic regulation of RNA splicing in human pancreatic islets": Methods

**Human pancreatic islet datasets**

We compiled 447 human islets samples, which comprised 89 samples from GEO accession number GSE50244 [1], 118 from EGA accession number EGAD00001001601 [2], 112 samples from T2DSystems [3] (EGAS00001005535), and 128 samples from CRG cohort (EGA). Out of 447 samples, 399 samples passed genotype and RNA-seq QC filters described below.

For the CRG cohort, human pancreatic islets from organ donors without a history of glucose intolerance were purified using established isolation procedures [4-7], shipped in culture medium and re-cultured at 37 °C in a humidified chamber with 5% CO2 in glucose-free RPMI 1640 supplemented with 10% fetal calf serum, 100 U ml−1 penicillin, 100 U ml−1 streptomycin and 11 mM glucose for 3 days before analysis. Islet isolation centers had permission to use islets for scientific research if they were insufficient for clinical transplantation following national regulations and ethical requirements and institutional approvals from University of Lille, University of Geneva, and Milano San Raffaele Hospital. Ethical approval for processing de-identified samples was granted by the Clinical Research Ethics Committee of Hospital Clinic de Barcelona and Parc de Recerca Biomèdica de Barcelona.

**Genotype processing**

The 128 CRG cohort samples were genotyped with Illumina Infinium OmniExpress 12 v1 and HumanOmni 2.5-8v1 on a total of 624K SNPs. The 89 samples from Fadista, J. et al 2014 (GSE50244) [1] were genotyped using Illumina HumanOmniExpress 12v1 C on a total of 609K SNPs. The 118 samples from Van de Bunt, M. et al. 2015 (EGAD00001001601) [2] were genotyped using Illumina Omni2.5+Exome array on a total of 2.6M SNPs. Finally, the 112 samples from T2DSystems (EGAS00001005535) [3] were genotyped using Illumina's Human Omni 2.5 exome array on a total of 2.6M SNPs.

A three-step quality control of genotype data, involving two stages of SNP removal and one intermediate stage of sample exclusion, was conducted in each cohort. Genotyped SNPs were filtered if (i) minor allele frequency (MAF) $< 0.01$), (ii) missing genotype rate $\geq$ 5% and (iii) significantly deviated from Hardy-Weinberg equilibrium (HWE p-value $\leq 1x10^{-6}$). Samples were excluded if (i) individual missing genotype rate $\geq$ 2%, (ii) cryptic relationships and sample duplicates (individuals with higher individual missingness genotype rate from pairs with pi $\geq 0.185$), or (iii) showed >4 standard deviations from the mean according to the first four principal components in each given cohort.

After QC analysis for genotype data (i) 113 individuals and 557,422 SNPs were retained for the CRG cohort, (ii) 112 individuals and 1,499,688 SNPs for the Van de Bunt, M. et al cohort [2] (iii) 89 individuals and 596,464 SNPs for the Fadista, J. et al, 2014 [1] cohort and (iv) 109 individuals and 1,543,968 SNPs for the T2DSystems cohort[3].

For each cohort, we generated per-chromosome VCF files after removing all strand ambiguous variants (AT or CG SNPs), and checking for strand alignment against the Haplotype Reference Consortium (HRC) and 1000 Genomes (1000G) reference SNP list. *HRC-1000G-check-bim.pl* script with the *-n* option (to turn off the removal of variants showing MAF differences between the reference panel and the study genotypes) was used. We submitted resulting VCF files to the Michigan Imputation Server (https://imputationserver.sph.umich.edu/index.html): EAGLE2 [8] was used for phasing, minimac3 [9] for genotype imputation with the HRC [10] r1.1and the 1000G Phase 3 release reference panels [11], independently. For each dataset of imputed genotypes, we excluded variants with: (i) MAF < 1%, (ii) imputation-quality $R^2 < 0.7$, and/or (iii) HWE p-value $\leq 1x10^{-6}$. We extracted indels from the 1000G Phase3 imputed results, filtered them using the aforementioned criteria and merged with the filtered HRC imputed dataset.

**RNA-Seq data alignment and QC**
Raw fastq files from all cohorts were aligned to hg19 genome build using STAR [12] using the options, *--outFilterMultimapNmax 1 --outSAMstrandField intronMotif --outSAMattributes All --twopassMode Basic*. WASP [13] pipeline was used to remove reads mapped with allelic bias. *VerifyBAMID* [14] was used to assess the concordance between genotypes and RNA-Seq and samples with more than 2% contamination (CHIPMIX > 0.02 and FREEMIX > 0.02) were removed. This resulted in 101 samples from CRG cohort, 109 from samples Van de Bunt, M. et al cohort [2], 82 samples from Fadista, J. et al, 2014 [1] cohort and 107 samples from T2DSystems [3]. The resulting BAM files were used for gene expression quantification and to calculate the splicing activity. In-case of EGAD00001001601 [2], only bam files were available; hence the initial STAR alignment step was not performed.

**Principal component analysis of genotypes**
To identify individuals of divergent ancestry and to characterize population structure of 399 human islet samples, we first selected a subset of genotyped SNPs that were common in all 4 cohorts, that also passed all our QC filters (see **Genotype processing**), and with MAF $\geq$ 1% and missingness < 5% across all the samples. We also excluded SNPs in high LD (pairwise $r^2 \leq 0.1$ within 1 Mb window), C/G and A/T SNPs to avoid strand mismatches, and those located in previously reported regions with long-range LD. We aggregated the 1000 Genomes Phase3 reference dataset using the set of overlapping variants. flashPCA [15] tool was used to calculate genetic principal components (PCs) (**Supplementary Figure 1c, d).** First four genetic PCs are used as covariates in subsequent sQTL and eQTL analysis.

**Gene expression quantification**
In-house developed human pancreatic islet transcriptome annotations built with a combination of CAGE as well as short and long sequence reads were used to quantify total gene expression (Goutham Atla and Anthony Beucher, unpublished). FeatureCounts [16] was used to obtain gene level qualifications using default parameters except using appropriate strandedness flag for each dataset. Genes with less than 5 raw reads mapped in less than 10% of the samples within each cohort were removed. CPM normalization was performed using edgeR [17] *cpm* function and then log2 transformed. Combat [18] was used to remove known sequencing batch effects. Principle components (PCs) were calculated on Combat corrected gene expression values using *prcomp* function in R (**Supplementary Figure 1a**).

**Splicing quantification**
To quantify splicing activity, we used the annotation free method, *leafcutter* [19]. Briefly *bam2junc.sh* script from leafcutter was used to quantify junction spanning reads based on spliced alignments from bam file. We removed junctions that are not supported by at least 5 spliced reads in at least 10% of samples. We then used *leafcutter_cluster.py* script with options *-m 30 -l 500000* to cluster the junctions anchoring on shared splice sites. This identifies local splicing events which are a cluster of alternatively used junctions. *prepare_phenotype_table.py* script was used to get relative junction usage (ratios) across samples. The relative junction usage is also referred to as percent spliced in (junction PSI) in the manuscript. Combat [18] was used to remove known sequencing batch effects and principal components were generated using *prcomp* function in R (**Supplementary Figure 1b**).

**cis-eQTL mapping**
*cis*-eQTL mapping was performed using QTLtools [20] for 399 samples with available QCed genotype and RNA-seq data using a cis-window of 500 kb up- and downstream of the

transcription start site (TSS). 15 PCs derived from gene expression and 4 genetic PCs were used as covariates in the linear model. In order to identify best associated cis eQTL SNP-eGene pairs, QTLtools was run using the permutation pass mode using default parameters and *--permute 1000 --window 500000 --seed 123456*. Beta approximated permutation p-values were adjusted for multiple testing correction using Storey q-values implemented in the *qvalue* R package. We set the significance threshold at FDR q-value ≤ 0.01. This resulted in 3433 genes (eGenes) with significant eQTLs (**Supplementary Table 1)**. We also calculated nominal p-values for all cis-SNPs within a 500kb window centered on the TSS of each gene (nominal pass mode from QTLtools, *--nominal 1 --window 500000 --seed 123456*). To identify all significant variant-gene pairs, we defined a genome-wide p-value threshold (*pt*), by considering the empirical p-value of the eGene closest to the 0.05 FDR threshold. A gene-based nominal p-value threshold was then calculated using *pt* and the beta distribution parameters from QTLtools. For each significant eGene, variants with a nominal p-value below the gene-level threshold were considered in subsequent analyses (significant nominal cis-eQTL variants) (**Supplementary Data 1**).

**cis-sQTL mapping**

We performed cis-sQTL mapping as described for cis-eQTL identification using intron excision ratios and a cis-window of 50 kb up- and downstream of the junction (*--window 50000*). In case of cis-sQTLs, 5 PCs derived from splicing ratios and 4 genetic PCs were used in the linear model. In order to identify best associated cis sQTL SNP-junction pairs, QTLtools was run using the permutation pass mode (1000 permutations), and beta approximated permutation p-values were adjusted for multiple test correction using Storey q-values implemented in the *qvalue* R package. We set the significance threshold at FDR q-value ≤ 0.01, resulting in 4,858 junctions with a significant sQTL (**Supplementary Table 2**) We also calculated nominal p-values for all cis-SNPs within a 50kb window around the junction (nominal pass mode from QTLtools) (**Supplementary Data 2**). To identify all significant variant-junction pairs, we defined a genome-wide p-value threshold (*pt*) by considering the empirical p-value of the junctions closest to the 0.05 FDR threshold. A junction-based nominal p-value threshold was then calculated using *pt* and the beta distribution parameters from QTLtools. For each significant junction, variants with a nominal p-value below the gene-level threshold were considered in subsequent analyses (significant nominal cis-eQTL variants).

**Annotation of sQTL junctions**

As leafcutter identifies junctions de-novo, we used transcriptome annotation GTF files as a base to annotate the junctions with respective genes. We used *gtf2leafcutter.pl* script from leafcutter's leafviz module (https://github.com/davidaknowles/leafcutter/tree/master/leafviz) to obtain the intron coordinates of all genes. The sQTL junctions were then mapped to the intron coordinates and annotated with respective gene names.

**Magnitude of genetic effects on splicing**

To quantify the magnitude of genetic effects on splicing, for each cluster we chose a junction with the best q-value. Then, for each sQTL junction, we calculated the difference in median junction usage (delta-psi) from samples with homozygous reference and homozygous alternate alleles and plotted as a function of the $-\log_{10}$ (q-value). If alternate homozygous samples were not available, we chose median junction usage from heterozygous samples.

**Visualization of splicing events**
The junctions identified from leafcutter were loaded into IGV [21] for visualization as arcs, and box plots were plotted using python.

**Comparison of eQTLs and sQTLs**
Our maps of genetic effects on splicing were compared to our eQTLs but also to previously reported exonQTLs and eQTLs in the largest study to date in human pancreatic islets by insPIRE consortium [22]. To estimate the gain of novel information uniquely provided by our sQTL analysis, we first determined the overlap between sGenes from sQTLs and eGenes from eQTLs in our study, and eQTLs and exon QTLs mapped by insPIRE [22]. For those genes that contain both sQTLs and either eQTLs or e/exon QTLs in insPIRE, we calculated the LD ($r^2$ measure) between the lead sQTL and all other SNPs within 1Mb using PLINK [23] (v1.9 *--ld-window-kb 1000 --ld-window 99999 --ld-window-r2 0)*. As a reference dataset for LD calculations, we used our high-quality genotypes in 399 samples. Then, for each gene we plot the LD $r^2$ distribution between the lead sQTL and the lead QTL from the dataset being compared, respectively.

**Genomic enrichment analysis of eQTLs and sQTLs**
We used GREGOR [24] to perform enrichment analysis of lead sQTLs and eQTLs in different genomic annotations using the following options `R2THRESHOLD=0.7, LDWINDOWSIZE=50000 (for sQTLs), LDWINDOWSIZE=100000 (for eQTLs), MIN_NEIGHBOR_NUM=200 and POPULATION=EUR.`

The human islet regulatory annotations were described previously [25]. For genic annotations, we used Gencode [26] v34 GTF file. We used *gtf2leafcutter.pl* script to obtain all exon, intron, 5' and 3' coordinates from GTF file. 5' and 3' coordinates were extended to +3bp into exon and +2bp into intron. All annotations were made sure to be contain mutually exclusive genomic space.

**Comparison with GTEx**
We obtained summary statistics data for eQTLs and sQTLs from 49 GTEx tissues [27]. We first listed eGenes and junctions with significant eQTLs and sQTLs, respectively, at FDR 0.01, consistent with our significance threshold. For the resulting significant variant-phenotype associations for each of the 49 tissues, variant and junction coordinates (for sQTLs) were lifted down from hg38 to hg19 using liftOver [28]. For each GTEx tissue, we looked at the variant-phenotype (eGenes or Junctions) overlap with our islet eQTL and sQTLs, using nominal QTL variants. For example, for GTEx *x* eGene in *j* tissue, if any of the GTEx significant variants mapped any of our nominal eQTL variants for that *x* eGene, we considered that islet eQTL signal to be shared with that given *j* tissue. Same approach was implemented to sQTLs. We excluded from this analysis testis, given the pervasive number of eQTLs, and pancreas because it is a partial surrogate of pancreatic islets.

**Quantile-quantile plots**

We generated quantile-quantile (Q-Q) plots to estimate genomic inflation in sQTLs and eQTLs for T2D risk, glycemic traits variation. For T2D we used BMI-adjusted T2D summary statistics [29]. For glycemic traits, we leveraged summary statistics data from a trans-ancestral meta-analysis for fasting glucose (FG) and fasting insulin (FI) [30]. We included variants with MAF ≥ 5% that were intersected with our nominal e- and sQTLs. For each trait comparison, we generated 1000 permutations of subsets of control sQTL variants to provide further support to the observed enrichment of e- and s-QTLs among GWAS variants. Each control set of sQTL-like variants was generated by first identifying independent LD blocks [31], that comprised nominal eQTL or sQTL variants, respectively. Then, we shuffled non-overlapping genomic regions, that were created based on the size of the genomic ranges where our nominal sQTL variants were located, across the genome, but excluding those independent LD regions where either eQTL or sQTL variants were located, blacklisted regions (wgEncodeDacMapabilityConsensusExcludable.bed.gz and wgEncodeDukeMapabilityRegionsExcludable.bed.gz) and the MHC region. Among the set of shuffled independent LD regions, we randomly sampled the same number of nominal sQTL variants. This was done 1000 times.

For the specific purpose of addressing T2D susceptibility across shared or independent sQTL/eQTL effects, we considered sQTLs and eQTLs from junctions and eGenes whose corresponding lead sQTL and eQTL were in $r^2 \geq 0.6$ as shared. The rest were considered as sQTL or eQTL specific genetic effects, respectively. To estimate T2D risk inflation across islet-selective sQTLs and eQTLs, we grouped sQTLs and eQTLs that were shared in ≤ 6 GTEx tissues (see **Comparison with GTEx** section) as islet-selective QTL effects. To assess the strength of T2D-risk inflation, we leveraged 1000 permutations of control-sets of sQTL-like variants described above.

**Colocalization analysis across T2D and FG/FI independent GWAS signals**

We collected 403 independent lead variants for T2D [29] and 277 for FG and FI [30]. For each trait, we performed colocalization as implemented in *gwas-pw* [32] at each independent GWAS signal. We only tested for colocalization with e and s-QTLs at those signals with at least one of credible set variant (genetic posterior probability ≥ 0.01) in LD ($r^2 \geq 0.6$) with a lead eQTL or sQTL, respectively. LD calculations were performed using the high-quality genotypes from our ~399 islet donor samples. If for a given independent GWAS signal, any of the available proxy variants was not included in our imputed genotypes, we used 1000 Genomes Phase3 genotypes (with European descent) as the reference panel [11]. Colocalization was performed across 1Mb genomic interval centred on the reported lead independent variant for a given GWAS locus. We nominated a region as a colocalized locus if the posterior probability for the model 3 (presence of the same genetic variant underlying the QTL and GWAS association, "colocalization") was ≥ 0.8. Colocalization signals were visualized using R-3.6.1 and LocusCompareR 1.0.0.

**Visualization of regional association plots**

Regional association plots for T2D and T1D susceptibility regions were created using LocusZoom [33] v1.4 in R-3.6.1.

**TWAS analysis**
FUSION [34] was used for TWAS analysis with T2D and glycemic traits (FG and FI). For gene expression, first the weights were computed using *FUSION.compute_weights.R* using options --*models top1, blup, bslmm, lasso, enet* on the same data that is used for eQTL analysis. 15 gene expression PCs and 4 genetic PCS were used as covariates and variants within a cis-window of 500kb from TSS were used. For splicing analysis, variants within 50kb from the junction boundaries were used and 5 splicing PCs and 4 genetic PCs were used to compute the weights.

After computing weights, *FUSION.assoc_test.R* script was used to test for association of the pre-computed weights and GWAS summary statistics data for T2D [29] and glycemic traits[30] (FG and FI). We only included GWAS data from variants that overlaid our ~6.5M high-quality imputed common genetic variants. Multiple test correction was applied to the resulting p-values using Bonferroni. We also excluded TWAS results with low colocalization posterior probabilities (gwas-pw PPA_3 or COLOC PP4 < 0.6) and with large confounding effects from linkage (COLOC PP3 > 0.8 and COLOC PP4 < COLOC PP3).

We mapped TWAS results to T2D and glycemic GWAS loci by identifying TWAS results whose best GWAS lead variant was in LD ($r^2 \geq 0.1$, calculated using the genotypes of individuals with European descent from the Phase 3 of the 1000 Genomes Project [11]) or less than 500kb away from lead GWAS signals, for each trait respectively. The rest of TWAS results were considered as novel GWAS loci. For T2D, we also considered as known GWAS loci where the TWAS signals is in LD ($r^2 \geq 0.1$) with lead independent signals identified in a recent large-scale meta-analysis [35].

**Single cell gene expression analysis**
We quantified published single cell data sets [36-39] against in-house annotations (Goutham Atla and Anthony Beucher, unpublished) using featureCounts [16]. Seurat V3 [40] was used to normalize and identify the cell-types within each data set. A gene is annotated as expressed if it has normalized counts of ≥ 0.3 in at least 30 cells of a given cell-type in one of the data sets.

**Network analysis**
We selected all the nominated effector genes for T2D, FG, FI from the current study and InsPIRE [22], filtered for 106 that encode for proteins with known or presumed function, and used them to construct networks using StringDB [41]. StringApp [42] in cytoscape [43] was used to identify the protein-protein interaction networks using default parameters, confidence score cutoff 0.4 and maximum additional interactors of 5. This yielded 1.8-fold higher protein-protein interactions than random sets, although only subnetworks with 3 or more components are shown.

**Credible set analysis**
We used fine-mapping approaches to identify candidate causal variants that underlie cis-eQTL and sQTL *loci* (**Supplementary Data 3 and 4, respectively)**. We identified 95% credible set variants using CAVIAR [44] software and allowing for one causal variant (*-c 1*). LD information between SNP pairs (i.e. the *r* matrix) was generated using PLINK[23] v1.9 –*matrix –r*, and as reference panel, we used our 399 high-quality human islet samples used in the eQTL and sQTL identification.
For each e/sQTL credible set, the posterior probabilities were plotted with respect to underlying genomic annotations.

**In-silico functional scores**

The potential impact on disease was assessed for the credible set variants of both eQTLs and sQTLs on the basis of their predicted transcriptional and post-transcriptional regulatory effects. We used a deep-leaning model that is trained on transcriptional regulatory features such as histone marks, DNAse I profiles and transcription factors, a total of 2,002 features, and is implemented as DeepSEA [45] as well as post-transcriptional regulatory features such as RNA-binding proteins binding data based on CLIP experiments on 82 unique RBPs (ENCODE and other CLIP datasets), which is implemented as Seqweaver [46]. We performed in-silico mutagenesis on both eQTL and sQTL credible set variants using both deepsea and seqweaver models and obtained the Disease Impact Scores (DIS) from respective models (**Supplementary Data 3 and 4, respectively**). Prior to in-silico mutagenesis, the strand information was added to sQTL credible sets based on the orientation of the target gene. The credible sets were further stratified based on their location in the genome and the DIS from both models for each category of variants were shown as boxplots.

**Integrating GWAS credible sets with QTL credible sets**

We obtained the pre-calculated genetic credible sets for T2D [29] (https://diagram-consortium.org/downloads.html). For each colocalized loci (colocalization PP >0.8), we then annotated credible set variants into two mutually exclusive categories. (i) A GWAS credible set variant which is also a QTL credible set variant. (ii) A GWAS credible set variant which is not a QTL credible set variant but in LD ($r^2 < 0.1$) with a lead QTL. We then plotted the distributions of CPPs of all GWAS credible sets stratified into above two categories. If a variant belongs to more than one credible set, we used the maximum CPP of that variant.
